# Critical roles of MCM8 in meiotic recombination during mouse spermatogenesis

**DOI:** 10.64898/2026.03.28.714908

**Authors:** Lava Kumar Surarapu, Kevin Tilton, Maria Rosaria Dello Stritto, Ananya Acharya, Andrea Marton Menendez, Min Lu, Najma Shaheen, Shun Liang, Mythri Iyer, Petr Cejka, Florencia Pratto, Devanshi Jain

**Author notes:** Division of Radiation and Genome Stability, Department of Radiation Oncology, Dana-Farber Cancer Institute, Harvard Medical School, Boston, MA, USA.

## Abstract

Meiotic DNA double-strand break (DSB) formation and repair by homologous recombination is crucial for ensuring proper chromosome segregation. In mice, the mini-chromosome maintenance family protein, MCM8, has been proposed to function in meiotic recombination and its loss leads to infertility, but the underlying mechanisms are poorly understood. Here we used cytological and genomic assays to infer the role of MCM8 during meiotic recombination in mouse spermatocytes. We show that MCM8-deficient spermatocytes exhibit increased levels of SPO11-dependent DSBs at recombination hotspots during early prophase. DSBs are resected normally and accumulate strand-exchange proteins. However, downstream recombination intermediates are barely detected and recombination intermediate-associated MutSgamma foci do not form efficiently. Consistent with a role in early recombination intermediate processing, MCM8 binds to displacement loop (D-loop) structures *in vitro*. We propose that MCM8 controls meiotic recombination in at least two ways. MCM8 participates in regulating meiotic DSB number. Further, MCM8 plays a role in the formation and/or stability of post-resection recombination intermediates, steps that are critical for DSB repair via recombination and for efficient synapsis of homologous chromosomes during mouse meiosis.

## Introduction

Homologous recombination during meiosis increases genetic diversity plus ensures homologous chromosome pairing and segregation in most species (Zickler and Kleckner 2015). Meiotic recombination initiates with the formation of a tightly controlled number of DNA double-strand-breaks (DSBs) (Lam and Keeney 2014). DSBs are nucleolytically processed (resected) creating single-stranded DNA (ssDNA) overhangs, which engage in homology search and perform strand exchange with the intact homologous chromosome, forming a recombination intermediate called the displacement loop (D-loop) (Brown and Bishop 2014; Hunter 2015). Extension of the overhang followed by ejection and annealing to the opposite DSB end repairs many DSBs as noncrossover events. At a subset of meiotic DSBs, the overhang is extended, and the D-loop engages the opposite DSB end, creating an intermediate that is most frequently resolved to yield reciprocal exchange of homolog arms called a crossover. Interhomolog interactions during recombination facilitate homolog pairing and crossovers between homologs ensure proper homolog segregation (Hunter 2015; Zickler and Kleckner 2015).

Members of the Mini-Chromosome Maintenance (MCM) protein subfamily of hexameric helicases have been implicated to be critical regulators of meiotic recombination (Blanton et al. 2005; Hartford et al. 2011; Lutzmann et al. 2012; Crismani et al. 2013). The MCM family includes the broadly conserved and essential replicative MCM helicase subunits (MCM2 through MCM7), and two lesser characterized members, MCM8 and MCM9 (Maiorano et al. 2006; Liu et al. 2009). *Mcm8* and *Mcm9* are widely conserved across eukaryotes, but absent in most fungal and *Caenorhabditis* lineages, and Schizophoran flies have *Mcm8* but not *Mcm9* (Liu et al. 2009; Kohl et al. 2012). In *Drosophila*, the MCM8 ortholog (REC) forms a complex with two MCM-like proteins, MEI-217 and MEI-218, and is required to form crossovers during meiosis (Blanton et al. 2005; Kohl et al. 2012). The loss of crossovers in *rec* mutants is suppressed by mutation of the Bloom syndrome helicase (BLM), which unwinds recombination intermediates that might otherwise be processed into crossovers, suggesting that MCM8 opposes this activity and stabilizes recombination intermediates to generate crossovers in flies. Contrastingly, *Arabidopsis* MCM8 (*At*MCM8) does not regulate crossover formation. *Atmcm8* mutants form normal numbers of crossovers but harbor fragmented chromosomes during anaphase, suggesting that *At*MCM8 plays a more general role in meiotic DSB repair (Crismani et al. 2013). Thus, the specific contributions of MCMs to meiotic recombination differ between species.

Mammalian MCM8 and MCM9 form a heterohexameric helicase that binds and unwinds branched DNA substrates *in vitro*, indicating that they function together (Li et al. 2021; McKinzey et al. 2023; Weng et al. 2023; Acharya et al. 2024). Indeed, in somatic cells MCM8 and MCM9 form a complex that has been reported to function in promoting replication fork progression and in homologous recombination-mediated DNA repair (Lutzmann et al. 2012; Nishimura et al. 2012; McKinzey et al. 2021; Griffin et al. 2022). Proposed functions during homologous recombination may be context dependent and include facilitating resection of breaks (Lee et al. 2015), recruitment of strand exchange proteins (Lutzmann et al. 2012; Park et al. 2013; McKinzey et al. 2021), as well as recombination-associated DNA synthesis (Natsume et al. 2017; Hustedt et al. 2019; Huang et al. 2020).

MCM8 and MCM9 have also been proposed to function during mammalian meiotic recombination, although their individual contributions differ (Hartford et al. 2011; Lutzmann et al. 2012). *Mcm9^-/-^* mice are defective in primordial germ cell proliferation which leads to reduced germ cells, surviving germ cells arrest during meiosis, but many cells complete meiosis and form sperm. Consequently, *Mcm9^-/-^*males are fertile. *Mcm8^-/-^* mice are also defective in primordial germ cell proliferation (Wen et al. 2024). *Mcm8^-/-^* germ cells additionally show meiotic defects consistent with a meiotic recombination defect, such as accumulation of recombination-associated factors and synaptic failure (Lutzmann et al. 2012). These defects result in meiotic arrest and *Mcm8^-/-^* males are infertile. How MCM8 facilitates meiotic recombination is unclear, however.

Here we utilize an *Mcm8* mutant allele isolated through a forward genetic screen to characterize the role of MCM8 in meiosis. We find that *Mcm8* mutants form higher than normal levels of meiotic DSBs. Additionally, while resection of meiotic DSBs appears to proceed normally, downstream repair and accompanying synapsis of homologs are severely compromised in mutants. We propose that MCM8 is required post-resection to form or stabilize recombination intermediates during meiosis.

## Results

### Isolation of an *Mcm8* mutant allele in a forward genetic screen for meiotic defects

To isolate mutant mice with meiotic defects, we performed a forward genetic screen (Jain et al. 2017; Jain et al. 2018; Petrillo et al. 2020; Guan et al. 2021; Sorkin et al. 2025). Briefly, we injected mice with *N*-ethyl-*N*-nitrosourea (ENU), which is an alkylating agent used to generate genome-wide point mutations (Caspary and Anderson 2006; Probst and Justice 2010). We next subjected treated mice to a three generation breeding scheme to homozygose the induced mutations (see Methods). Finally, we screened third-generation males for meiotic defects by immunostaining squashed preparations of testis cells for markers that report on meiotic prophase I progression.

We examined immunostaining patterns of SYCP3, which is a component of chromosome axes during meiotic prophase (Lammers et al. 1994; Zickler and Kleckner 2023). SYCP3-positive axial elements begin to form during the leptotene stage of meiotic prophase (**Figure 1A**). Axes elongate during zygotene, and homologous axes begin to synapse forming stretches of the synaptonemal complex (SC), a tripartite structure comprised of axial elements and the central region connecting them. Homologous axes are connected by the SC along their entire lengths during pachytene, and the SC disassembles during diplotene. We also tracked meiotic recombination by staining for γH2AX, which is the phosphorylated form of histone H2AX and forms in response to double-strand breaks (DSBs) (Mahadevaiah et al. 2001). γH2AX staining appears pan-nuclear during leptotene and zygotene when meiotic DSBs are forming (**Figure 1A**). During pachynema and diplonema, γH2AX largely disappears from autosomes as recombination proceeds, and recombination-independent γH2AX forms on the silenced X and Y chromosomes in the sex body.

**Figure 1.**
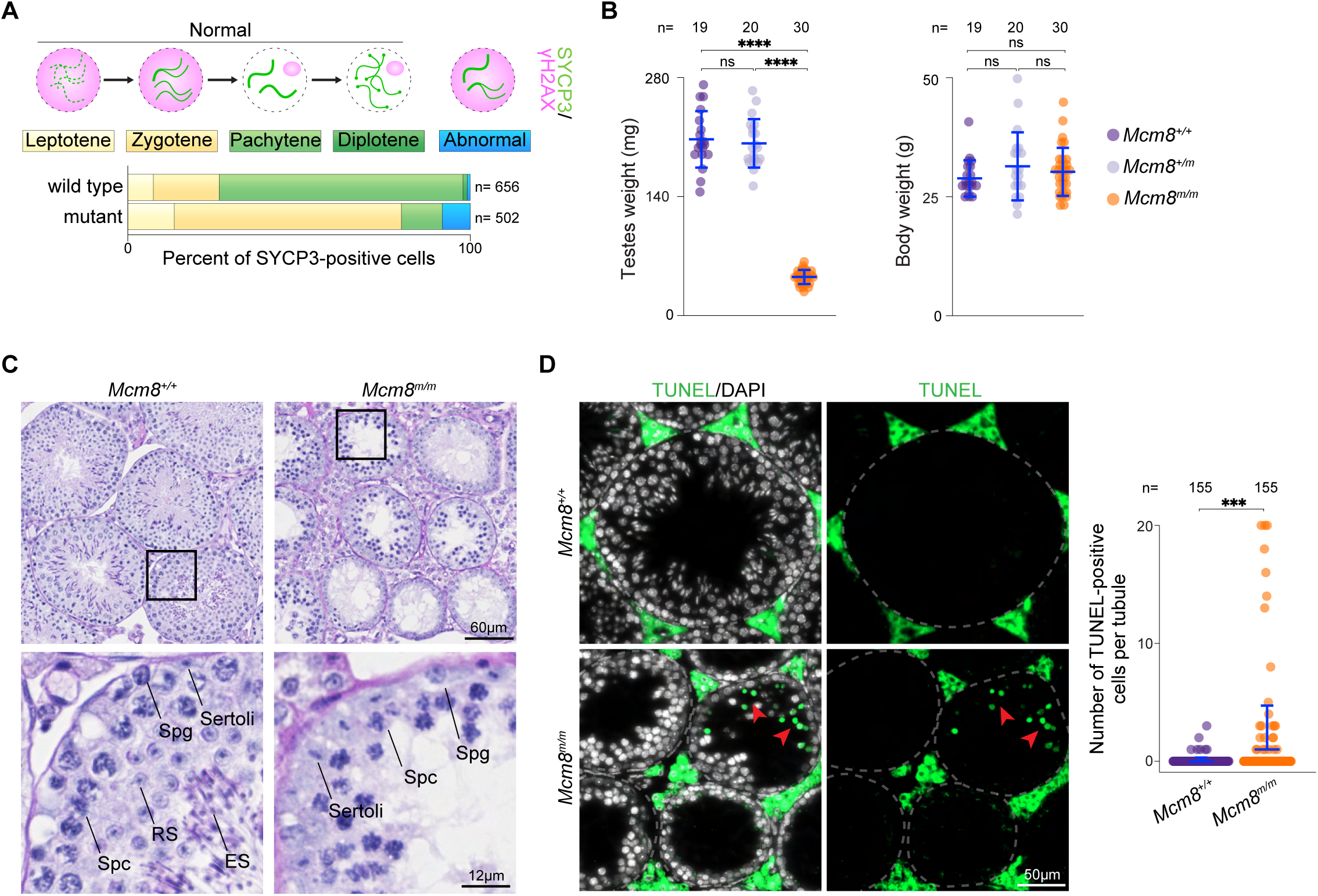
Males from the ENU-induced mutant line *mara* (*Mcm8^m^*) undergo abnormal meiosis. **(A)** Schematic depicting expected immunostaining patterns of SYCP3 and γH2AX in spermatocyte nuclei during leptotene, zygotene, pachytene and diplotene stages of meiotic prophase I, along with an abnormal staining pattern (pan-nuclear γH2AX with pachytene-like tracks of SYCP3). Distribution of meiotic prophase stages in a mutant mouse and a wild-type littermate from the *mara* line is shown below. Meiotic prophase stages were assessed using SYCP3 and γH2AX-staining patterns in squashed spermatocyte nuclei. (**B**) Testes (left) and body (right) weights of adult *Mcm8^+/+^*, *Mcm8^+/m^*, and *Mcm8^m/m^* mice. (**C**) Periodic acid-Schiff (PAS)-stained sections of Bouin’s-fixed testes from adult *Mcm8^m/m^*and *Mcm8^+/+^* littermates. Zoomed-in views of the regions indicated by black boxes are shown below and the following cell types are annotated: Sertoli, spermatogonia (Spg), spermatocytes (Spc), round spermatids (RS) and elongated spermatids (ES). (**D**) Images of TUNEL assay on testis sections from adult *Mcm8^m/m^* and *Mcm8^+/+^* littermates (left) and quantification of TUNEL assay (right). Red arrowheads within images point to TUNEL-positive cells (green). In panels B and D, blue lines are means ± standard deviations and results of two-tailed t tests are indicated (ns denotes not significant (p > 0.05), **** is p ≤ 0.0001; *** is p ≤ 0.001).

Through this screen we isolated a mutant mouse line that displayed SYCP3 and γH2AX staining patterns indicative of abnormal meiotic prophase progression (**Figure 1A**). We named this mouse line *mara* (**<u>m</u>**eiosis **<u>a</u>**nd **<u>r</u>**ecombination **<u>a</u>**ffected). *Mara* mutants accumulated spermatocytes with SYCP3 staining indicative of leptotene and zygotene stages, however cells with pachytene- and diplotene-like SYCP3 staining were depleted compared to normal littermates. Additionally, *mara* mutant spermatocytes acquired γH2AX staining in leptotene- and zygotene-like cells, but also contained abnormal cells with γH2AX present alongside tracks of SYCP3 staining consistent with levels of synapsis that are normally present in later prophase stages (**Figure 1A**). These staining patterns resemble those seen in meiotic mutants that are unable to complete synapsis and meiotic recombination (Mahadevaiah et al. 2001), like *Msh5^-/-^*(de Vries et al. 1999; Edelmann et al. 1999) and *Dmc1^-/-^* (Pittman et al. 1998; Yoshida et al. 1998). We identified a single nucleotide substitution within the *Mcm8* gene (*Mcm8^m^*) and *Mcm8*-null mice have been previously shown to be defective at meiosis, therefore we surmised that the *Mcm8^m^* mutation is the likely causal mutation for the *mara* phenotype.

*Mcm8^m/m^* mice had significantly reduced testes weights compared to wild-type and heterozygous littermates (78% and 77% reduction, respectively; **Figure 1B**), and *Mcm8^m/m^* body weights were similar to controls (**Figure 1B**). In histological analyses of testes, control animals showed populated seminiferous tubules with the full array of spermatogenic cell types including spermatids, while *Mcm8^m/m^* harboured shrunken seminiferous tubules lacking post-meiotic spermatids (**Figure 1C**). Some *Mcm8^m/m^*tubules contained primary spermatocytes along with Sertoli cells and spermatogonia, and some tubules appeared to be more severely depleted. Consistent with this, *Mcm8^m/m^* seminiferous tubules contained a significantly higher number of TUNEL-positive cells compared to control littermates, indicative of apoptosis (**Figure 1D**). These phenotypes are expected to result in sterility. While heterozygotes had normal fertility and Mendelian transmission of the *mara* mutation (29.34% *Mcm8^+/+^*, 53.10% *Mcm8^+/m^*, and 17.56% *Mcm8^m/m^* from heterozygote × heterozygote crosses; n = 467 mice; p=0.52, Fisher’s exact test), none of the three *Mcm8^m/m^* males paired with wild-type females for 12 weeks sired offspring.

### *Mcm8^m^* mutation causes loss of MCM8 protein

The hypogonadism and infertility phenotypes observed in *Mcm8*-*mara* mutants resemble those previously described for *Mcm8*^-/-^ males that lack detectable MCM8 protein in testis (Lutzmann et al. 2012), so we evaluated the impact of the *mara* mutation on *Mcm8* expression. The *mara* variant is a T to A nucleotide transversion at position Chr2:132,828,750 within the 10^th^ intron of *Mcm8* (**Figure 2A**). Examination of *Mcm8* mRNA by reverse-transcription PCR (RT-PCR) using primers spanning intron 10 yielded a PCR fragment of the expected length (103 bp) and an additional longer fragment (212 bp) that was absent in littermate controls (**Figure 2A**, **B**). Sanger sequencing the longer fragment revealed an insertion of a 109-bp-long stretch of intronic sequence located adjacent to the *mara* mutation. The A to T *mara* mutation creates a donor splice site consensus sequence (GTAAGT/GUAAGU) (Malard et al. 2022) that may be recognized by the splicing machinery, leading to aberrant splicing and inclusion of intronic sequence in *Mcm8^m^* mRNA.

**Figure 2.**
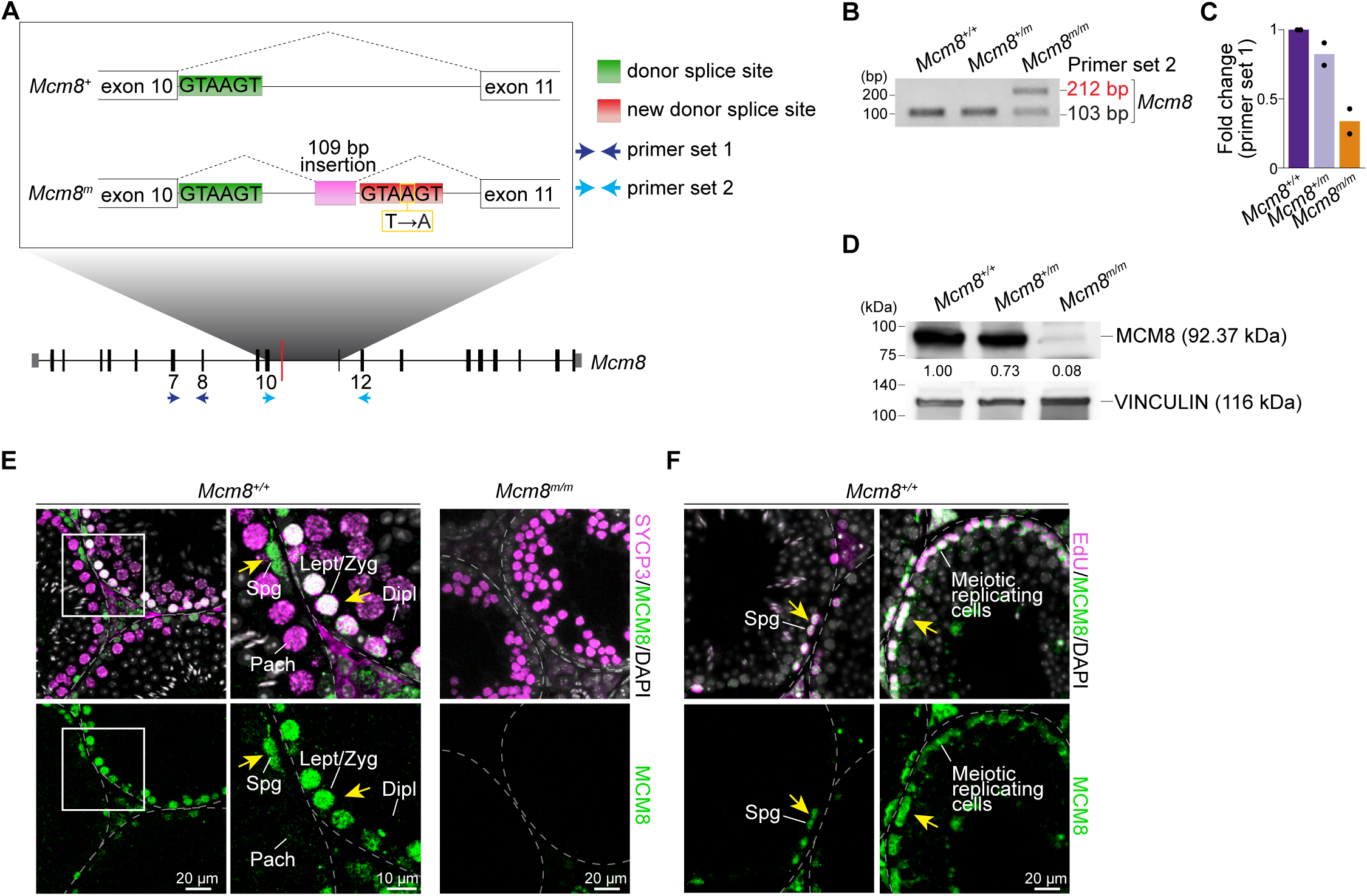
Characterization of the *Mcm8^m^* mutation and localization of MCM8 in testis. **(A)** Schematic describing the *Mcm8^m^*allele. Top: zoomed-in view of the intronic region containing the ENU-induced *mara* mutation (yellow box; T changed to A). The wild-type donor splice site (green) and a presumptive new donor splice site (red) created by the *mara* mutation, along with the path of splicing events (dotted lines) are annotated. The *mara* mutation leads to a 109-bp long insertion (pink) of intronic sequence in *Mcm8^m^* mRNA. Bottom: *Mcm8* exon map. Primer sets 1 (dark blue arrows) and 2 (light blue arrows) were used for RT-PCR in (**C**) and (**B**), respectively. (**B**) RT-PCR analysis of *Mcm8* in whole-testis RNA samples from adult mice of the indicated genotypes. PCR amplification was done using primers flanking the *mara* mutation-containing intron (primer set 2, shown in light blue in Figure 2A) and analyzed by gel electrophoresis. (**C**) Quantitative RT-PCR analysis of *Mcm8* in whole-testis RNA samples from adult mice using primers that are upstream of the *mara* mutation (primer set 1, shown in dark blue in Figure 2A; normalized to *Actb*). (**D**) MCM8 and VINCULIN (loading control) immunoblot in whole-testis extracts from adult animals. Relative MCM8 signal intensities normalized to VINCULIN are indicated. (**E**) Immunofluorescence of MCM8 and SYCP3 on adult testis sections. Regions indicated by white boxes in wild-type images are shown at a higher magnification to the right. Yellow arrows point to MCM8-positive cells. Spermatogonia (Spg), leptotene/zygotene (Lept/Zyg), pachytene (Pach), and diplotene (Dipl) cells are indicated. (**F**) Immunofluorescence of MCM8 and EdU on adult testis sections. Spermatogonia (Spg) and meiotic replicating cells are labeled. Yellow arrows point to cells that are MCM8-positive.

Mis-splicing and inclusion of a 109-bp-long intronic segment in *Mcm8* mRNA is expected to result in a frameshift, nonsense-mediated decay and/or premature termination of translation yielding a shorter, likely non-functional peptide with an alternative amino-acid sequence. Quantitative RT-PCR using primers spanning a segment of *Mcm8* that is upstream of the *mara* mutation showed reduced amplification in testes from *Mcm8^m/m^* compared to control littermates (**Figure 2A**, **C, Supplementary Figure 1A**), suggesting that *Mcm8* mRNA level is decreased in mutants. We next examined MCM8 protein levels by immunoblotting testis extracts. MCM8 protein level was slightly reduced in *Mcm8^+/m^*heterozygotes (∼27%) and more substantially reduced in *Mcm8^m/m^*(∼90%) compared to wild-type littermates (**Figure 2D, Supplementary Figure 1B**). Although lack of a suitable MCM8 antibody that recognizes the N-terminus precludes analyses of whether a truncated MCM8 protein is produced in *Mcm8^m/m^*, we surmise that a truncated protein is unlikely to be functional as the truncation is expected to result in loss of the C-terminal AAA+ ATPase domain. We conclude that the *mara* mutation culminates in loss of full-length MCM8 protein and that the *Mcm8^m^* allele is a null-like or severe loss of function allele.

To assess the temporal and spatial distribution of MCM8 during spermatogenesis, we immunostained testis sections from adult animals. MCM8 staining was prominent in SYCP3-positive spermatocytes in wild type (**Figure 2E**), and was undetectable in *Mcm8^m/m^* testes, consistent with the lack of immunoblot signal (**Figure 2D, Supplementary Figure 1B**). In wild-type spermatocytes, strong nuclear staining was present in leptotene and zygotene cells, and little to no staining was detected in pachytene and diplotene cells in sections. In addition to spermatocytes, we also observed MCM8 staining in spermatogonia (**Figure 2E**). MCM8 has been reported to promote replication fork progression in cell lines (Griffin et al. 2022), so we evaluated whether MCM8 is present in replicating cells during spermatogenesis. We subjected wild-type mice to a 2-hr pulse label with EdU *in vivo*, and subsequently immunostained testis sections. EdU-positive cells, inferred to be replicating spermatogonia and cells that have undergone meiotic replication, were frequently positive for nuclear MCM8 staining (**Figure 2F**).

### Defective spermatogenesis in *Mcm8^m/m^* mutants

MCM8 localisation patterns in meiotic and pre-meiotic cells and the abnormal meiotic progression observed in *mara* mutants during screening led us to evaluate the effects of MCM8 loss on spermatogenesis more closely. Testis sections from *Mcm8^m/m^* and wild-type or *Mcm8^+/m^* control littermates were immunostained for SYCP3 and γH2AX and the cellular composition of seminiferous tubules was evaluated (**Figure 3A**). As expected, control tubules contained either a double layer of spermatocytes (35.0% of tubules), comprised of one layer containing leptotene or zygotene cells and the second containing pachytene and sometimes diplotene cells, or tubules contained a single layer of pachytene cells (65.0% of tubules). In contrast, 37.5% of *Mcm8^m/m^*tubules were devoid of SYCP3-positive cells and most SYCP3-positive tubules contained only leptotene- or zygotene-like cells (57.0% of all tubules). Only 5.5% of *Mcm8^m/m^* seminiferous tubules had cells with pachytene-like SYCP3 staining (0.5% of tubules contained only pachytene-like cells and 5.0% of tubules contained leptotene- or zygotene-like along with pachytene-like cells). These patterns indicate that *Mcm8^m/m^* spermatocytes are able to proceed through leptotene and zygotene, and that spermatocytes arrest and/or die at or just prior to pachytene. Consistent with this, the numbers of leptotene- and zygotene-like cells within *Mcm8^m/m^* tubules was comparable or elevated compared to littermate controls (**Figure 3B**), but the numbers of pachytene-like cells were substantially lower (**Figure 3C**; 0–3 cells per tubule in mutants and ∼30–70 cells in wild-type or heterozygous littermates).

**Figure 3.**
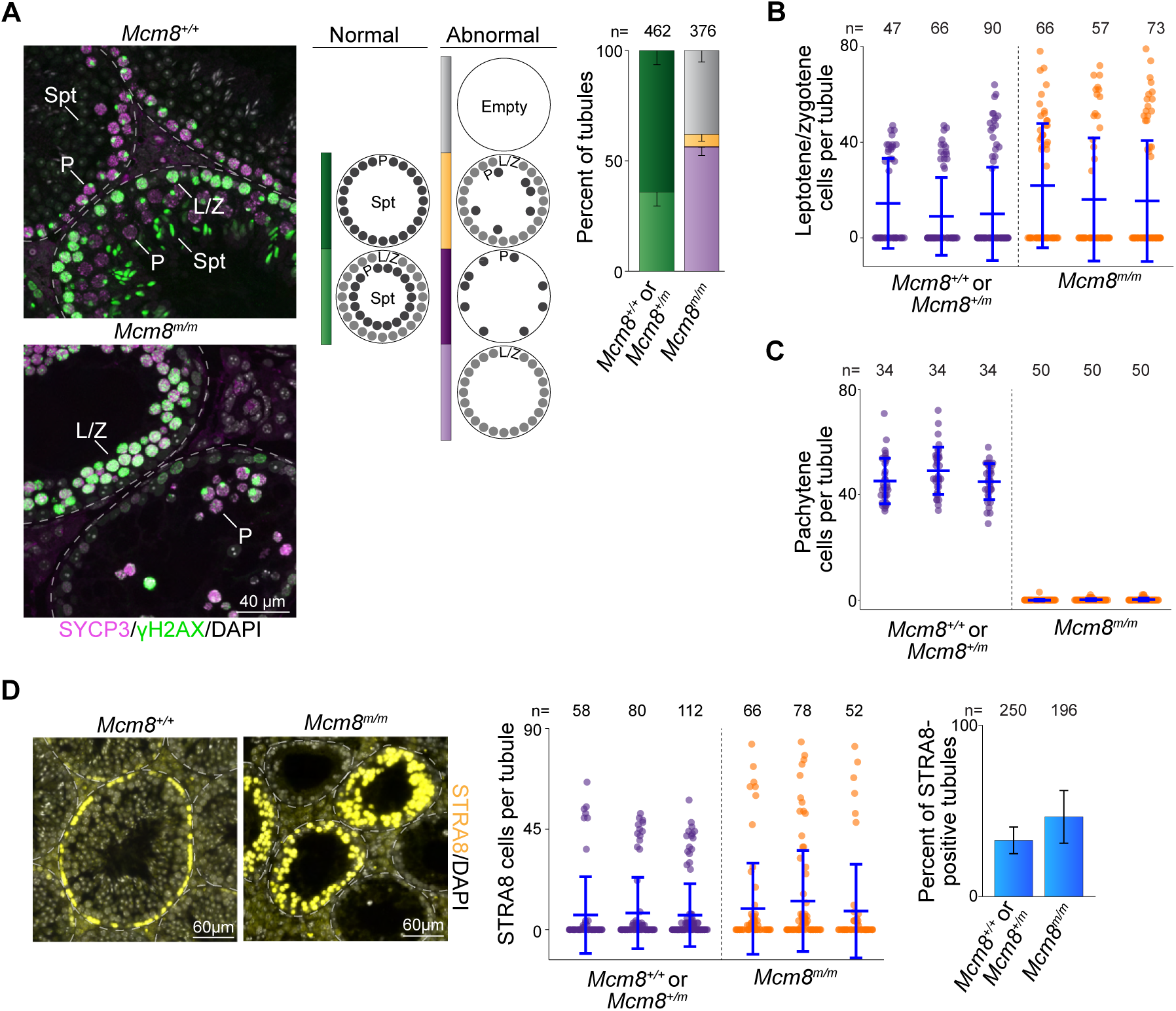
MCM8 deficiency perturbs meiotic progression. **(A)** Representative images of SYCP3- and γH2AX-stained adult testis sections (left), along with distribution of seminiferous tubule types (right). Tubule types were classified based on the presence of meiotic prophase cells, as judged by SYCP3- and γH2AX-staining patterns. Normal (wild type) tubule types and abnormal tubule categories observed in mutants are diagrammed. Leptotene/zygotene cells (L/Z), pachytene cells (P), and spermatids (Spt) are indicated within images. (**B**) Quantification of leptotene or zygotene cells in tubules. (**C**) Quantification of pachytene cells in tubules. (**D**) Images of STRA8-stained adult testis sections (left), along with quantification of STRA8-positive cells in tubules (middle and right). Bar plot shows means and standard deviations for three mice. In **B**, **C** and **D**, dot plots show numbers of positive-staining cells in individual tubule sections, with means ± standard deviations depicted using blue lines.

Because we observed MCM8 staining in meiotic replicating cells (**Figure 2F**), we evaluated this stage in mutants. We immunostained testis sections for STRA8, which is expressed in response to retinoic acid during preleptotene when cells are undergoing meiotic replication, and transiently expressed in differentiating spermatogonia (Oulad-Abdelghani et al. 1996; Koubova et al. 2006; Zhou et al. 2008). In testis sections from *Mcm8^+/+^* or *Mcm8^+/m^*, STRA8 staining showed a typical bimodal distribution (**Figure 3D**). Tubules contained either very few or no STRA8-positive cells, or contained a full layer of STRA8-positive cells that represents the meiotic replicating population (30–60 cells per tubule). *Mcm8^m/m^* testis sections showed a more dispersed distribution (**Figure 3D**). Many tubules contained few STRA8-positive cells. Other *Mcm8^m/m^* tubules contained higher numbers of STRA8-positive cells (∼30–90 cells per tubule), which we infer to be the meiotic replicating population. The numbers of STRA8-positive cells per tubule in this category were sometimes higher than those in control littermates, suggesting an increase in the numbers of cells undergoing meiotic replication. We expect that a prolonged meiotic replication stage in mutants would result in a higher number of tubules containing meiotic replicating cells, but this was not seen. The proportion of tubules that stained positive for STRA8 was comparable between mutants and control littermates (47% in *Mcm8^m/m^* and 33% in controls; **Figure 3D**). We therefore attribute the variation in the distribution of STRA8-positive cells in mutants to an increase in the numbers of cells entering meiosis and therefore undergoing meiotic replication. This interpretation is consistent with the observed increase in the numbers of leptotene and zygotene cells within mutant seminiferous tubules (**Figure 3B**).

MCM8 has been previously reported to be important for primordial germ cell proliferation (Wen et al. 2024), so we examined meiotic prophase in juvenile males at a developmental period when spermatocytes first appear and undergo meiosis (Bellve et al. 1977). Wild-type or *Mcm8^+/m^* testis sections immunostained for SYCP3 and γH2AX showed staining patterns indicative of normal meiotic progression (**Supplementary Figure 2A**, **B**). 60.0% of seminiferous tubules in testis sections from 13 dpp mice contained meiotic cells. By 16 dpp, almost all control tubules (97.0%) were populated with meiotic cells. The appearance of meiotic cells in *Mcm8^m/m^* was delayed, however. Meiotic cells were present in 19.0% of *Mcm8^m/m^* tubules at 13 dpp. At 16 dpp, 60.0% of *Mcm8^m/m^* tubules contained meiotic cells, while 40.0% remained devoid of SYCP3 staining. A straightforward interpretation of these staining patterns is that the depletion of primordial germ cells in MCM8-deficient animals (Wen et al. 2024) results in fewer spermatocytes during early development. And that in adult mice, once multiple waves of spermatogenesis have occurred, spermatogonial numbers may recover to yield higher numbers of spermatocytes entering meiosis (**Figure 3B**, **D**). Collectively, our results suggest that MCM8-deficient spermatocytes enter meiosis but undergo spermatogenic arrest during meiotic prophase leading to depleted seminiferous tubules, consistent with previous analyses (Lutzmann et al. 2012).

### Synaptic defects in *Mcm8^m/m^* meiosis

Defects in meiotic recombination and synapsis cause spermatocyte apoptosis during meiotic prophase (de Rooij and de Boer 2003) and have been previously observed in *Mcm8*-null mice (Lutzmann et al. 2012). Therefore, we examined synapsis and recombination in *Mcm8^m/m^*spermatocytes in detail.

We tracked synapsis by immunostaining spread spermatocyte nuclei for the chromosome axis protein SYCP3 and for the SC central region protein SYCP1 (de Vries et al. 2005). While SYCP3-positive chromosome axes were readily formed and synapsis appeared to be initiated in *Mcm8^m/m^*, most spermatocytes failed to achieve complete synapsis. In adult wild type, 66% of spermatocytes with elongated SYCP3 axes showed complete autosomal synapsis and 34% had partially synapsed chromosomes (**Figure 4A**). In contrast, 91% of *Mcm8^m/m^* spermatocytes had chromosomes with elongated SYCP3 axes that were juxtaposed and contained SYCP1 staining along varying stretches of the axes’ lengths (partial synapsis in **Figure 4A**) and 6% contained unaligned and unsynapsed axes with no SYCP1 staining (no synapsis in **Figure 4A**). Only 3% of spermatocytes appeared to have completely synapsed autosomes with SYCP3 and SYCP1 co-staining along entire axes lengths (full synapsis in **Figure 4A**), consistent with the depletion of pachytene-like cells in sections (**Figure 3A**, **C**).

**Figure 4.**
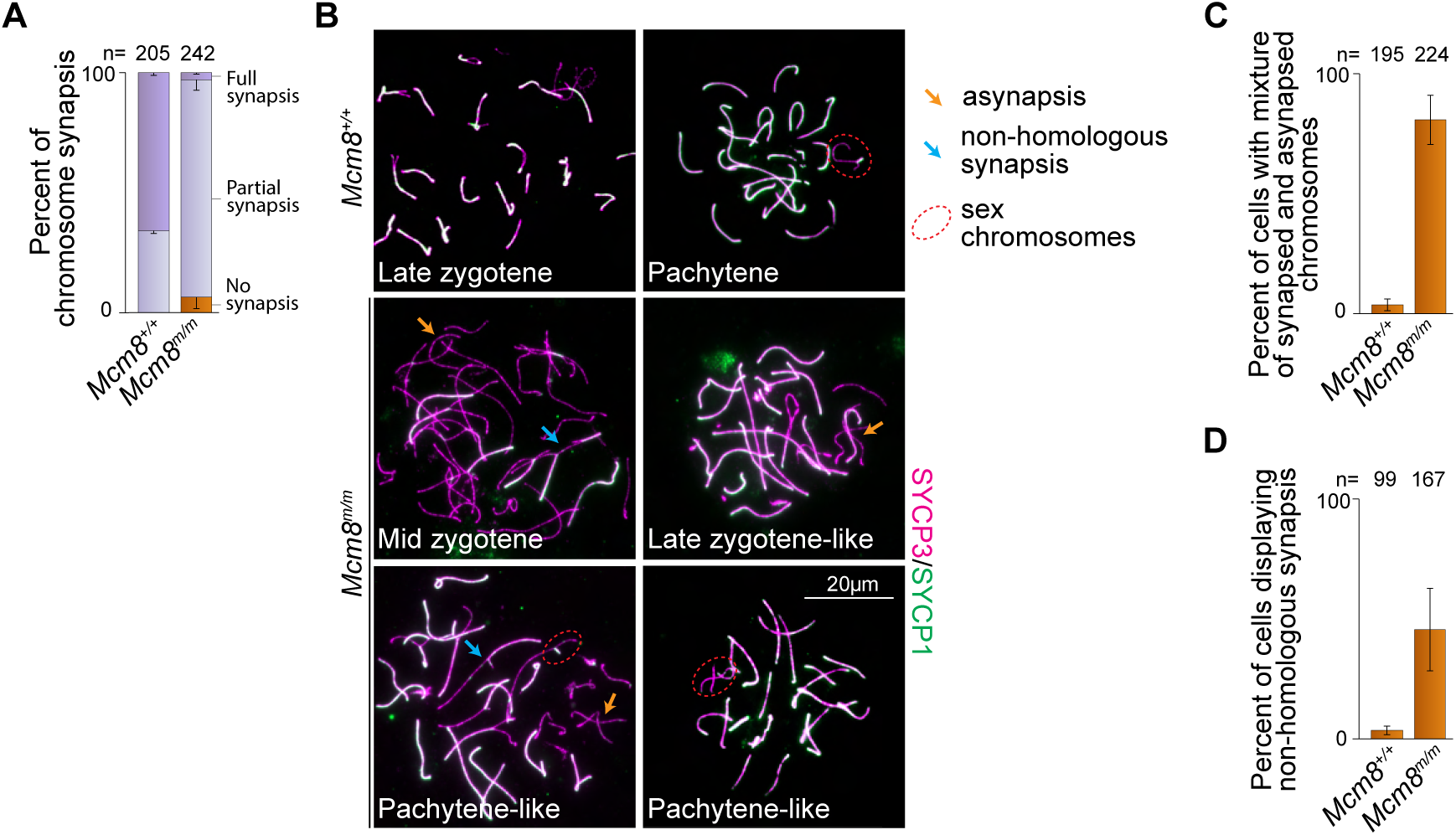
Synaptic defects in mice lacking MCM8. **(A)** Quantification of synaptonemal complex formation. Spermatocytes with elongated SYCP3 axes were scored and categorized as no synapsis (SYCP1 absent), partial synapsis (one or more stretches of SYCP1 present along juxtaposed SYCP3 axes) or full synapsis (SYCP1 present along all SYCP3 axes on autosomes). (**B**) Chromosome spreads immunostained for SYCP3 and SYCP1. Synapsis defects observed in *Mcm8^m/m^* such as asynapsed chromosomes (orange arrows) present alongside completely synapsed chromosomes and chromosomes displaying nonhomologous synapsis (blue arrows) are indicated. Sex chromosomes (red, dotted ovals) are highlighted in late stage cells. (**C**) Quantification of autosomal asynapsis. Spermatocytes with elongated SYCP3 axes and at least one fully-synapsed chromosome (SYCP1 present along the entire length of paired SYCP3 axes) were scored. (**D**) Quantification of non-homologous synapsis. Spermatocytes with elongated SYCP3 axes were scored. Bar plots show means and standard deviations for three mice.

In subsequent chromosome spread analyses, we refer to *Mcm8^m/m^*spermatocytes with partially synapsed autosomes as zygotene (**Figure 4B**), and further subcategorize based on increasing extents of autosomal synapsis as follows: early zygotene (little to no synapsis), mid zygotene (some synapsis) and late zygotene-like (extensive synapsis). Autosomal synapsis is succeeded by X-Y synapsis, occurring during pachytene in normal meiosis (Kauppi et al. 2011; Kauppi et al. 2012). We refer to *Mcm8^m/m^* spermatocytes in which sex chromosomes appear synapsed alongside extensive autosomal synapsis as pachytene-like (**Figure 4B**). Mutant spermatocytes in which sex chromosomes remain unsynapsed but autosomes appear completely synapsed are also categorized as pachytene-like.

In addition to the failure to achieve full synapsis, *Mcm8^m/m^*spermatocytes displayed synaptic abnormalities that are rarely observed in wild type. 81% of *Mcm8^m/m^* zygotene and pachytene-like spermatocytes harboured asynaptic chromosomes present alongside fully-synapsed chromosomes (orange arrows in **Figure 4B**, **Figure 4C**), while 4% of wild-type zygotene/pachytene cells displayed this abnormality. In normal zygotene, SYCP3 axes elongate at the same time that homologous synapsis ensues across all autosomes, therefore the presence of asynaptic chromosomes in cells where SYCP3 axes are elongated, and few chromosomes have completed synapsis indicates a defect in synapsis progression. Additionally, 46% of *Mcm8^m/m^* zygotene and pachytene-like cells contained a combination of unsynapsed and synapsed axes with partner switches indicative of nonhomologous synapsis (blue arrows in **Figure 4B**, **Figure 4D**), while nonhomologous synapsis was detected in 4% of wild-type zygotene/pachytene spermatocytes. Persistently asynapsed chromosome axes can undergo nonhomologous synapses (Kauppi et al. 2013), so this behaviour may stem from the inability of *Mcm8^m/m^*spermatocytes to undergo timely homologous synapsis. We conclude that MCM8 is critical for efficient synapsis of homologous chromosomes during meiosis.

### Recombination defects in *Mcm8^m/m^* meiosis

Our initial screening results indicated that *Mcm8^m/m^*mice contain abnormal spermatocytes with γH2AX present alongside seemingly synapsed chromosomes (**Figure 1A**). γH2AX forms in response to meiotic DSBs, progressively disappears as synapsis and recombination proceed, and is largely gone from synapsed autosomes in pachytene (Mahadevaiah et al. 2001), so the presence of γH2AX on synapsed chromosomes indicates the presence of persistent DSBs. We reexamined γH2AX staining, this time on spread spermatocyte chromosomes to better resolve staining patterns and their correlations with synapsis state. In most *Mcm8^+/+^* pachytene cells, γH2AX staining was largely gone from synapsed autosomes and restricted to the sex chromosomes (**Figure 5A**), as expected (Mahadevaiah et al. 2001). However, *Mcm8^m/m^*pachytene-like spermatocytes frequently displayed flares and/or patches of γH2AX on synapsed autosomes in addition to sex chromosome-associated γH2AX (**Figure 5A**), suggesting that meiotic DSB repair is dysregulated.

**Figure 5.**
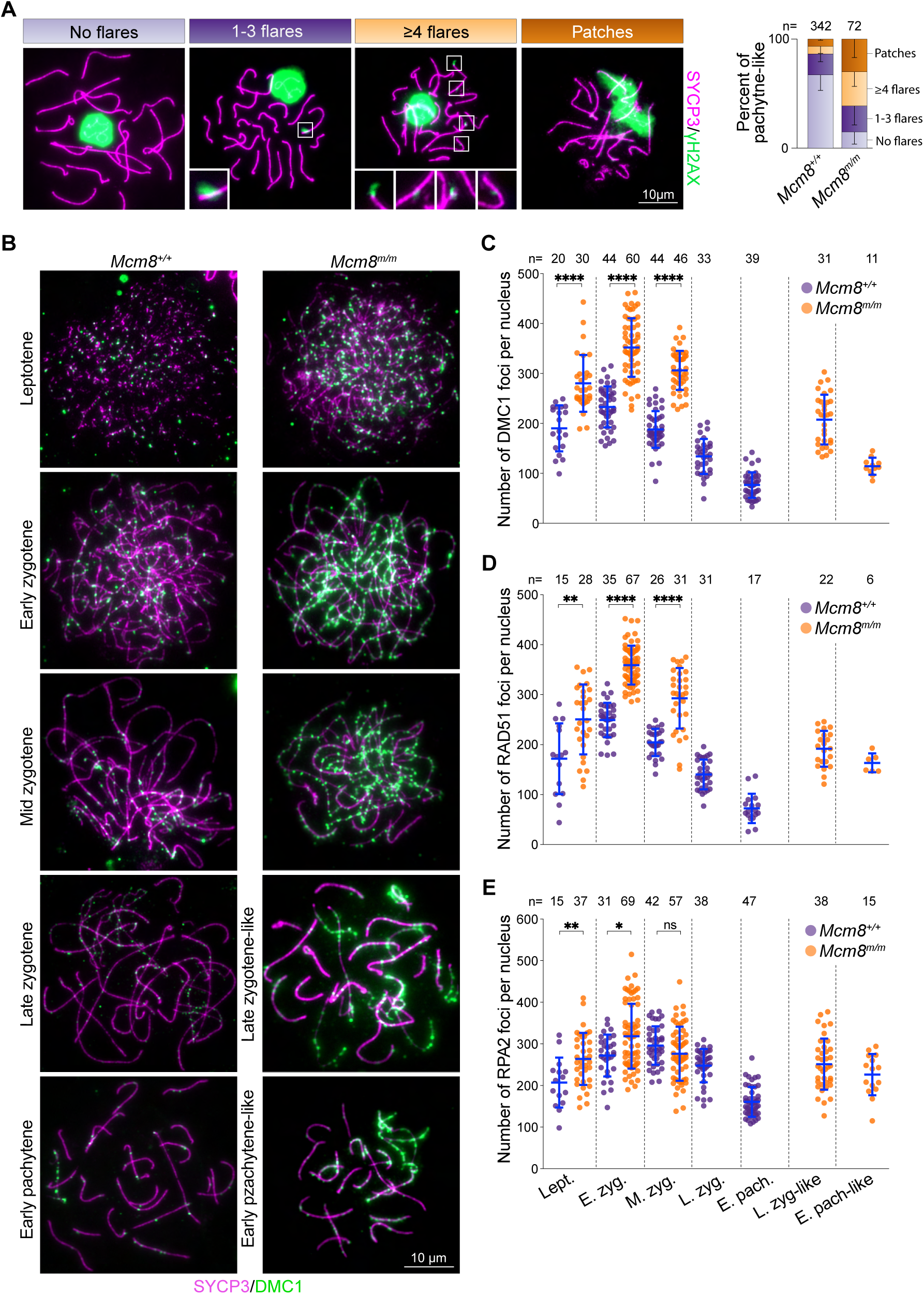
Recombination defects in mice lacking MCM8. **(A)** Chromosome spreads of *Mcm8^m/m^* pachytene-like cells with normal γH2AX staining (no flares), γH2AX flares, or γH2AX patches are shown on the left. Insets show higher magnification views of flares. Graph quantifies γH2AX staining patterns. Bars are means with standard deviations indicated for three mice. (**B**) Chromosome spreads depicting time course of DMC1 staining during meiotic prophase. (**C**) Quantification of DMC1 foci across meiotic prophase. (**D**) Quantification of RAD51 foci across meiotic prophase. (**E**) Quantification of RPA2 foci across meiotic prophase. In dot plots, foci overlapping with SYCP3-positive axes were scored. Each point is a count from one cell and means ± standard deviations for three mice are shown in blue lines. Results of two-tailed Mann Whitney tests are shown: ns is non-significant (p > 0.05), * is p ≤ 0.05; ** is p ≤ 0.01, *** p ≤ 0.001, and **** is p ≤ 0.0001. Cells were staged based on SYCP3-staining patterns: leptotene (Lept.), early zygotene (E. Zyg.), mid zygotene (M. Zyg.), late zygotene (L. Zyg.), early pachytene (E. Pach.), late zygotene-like (L. Zyg-like), early pachytene-like (E. Pach-like).

To assess the progression of meiotic recombination, we immunostained chromosomes for DMC1, RAD51 and RPA2 (**Figure 5B–E, Supplementary Figure 3**). DMC1 and RAD51 are strand-exchange proteins and orthologs of bacterial RecA (Brown and Bishop 2014). RPA2 is a subunit of the heterotrimeric single-strand DNA (ssDNA) binding protein RPA (Wold 1997). ssDNA formed at meiotic DSB sites is initially bound by RPA, which is subsequently replaced by DMC1 and RAD51 (Moens et al. 2002; Brown and Bishop 2014; Hinch et al. 2020). RPA also accumulates on ssDNA-containing recombination intermediates, such as D-loops.

In normal meiosis, chromosome axes-associated foci of DMC1 and RAD51 appear in leptotene, accumulate to maximal levels in early zygotene, and decline as DSB repair proceeds through pachytene, persisting longer on unsynapsed regions of the sex chromosomes (**Figure 5B–D, Supplementary Figure 3A**). In *Mcm8^m/m^*leptotene spermatocytes, DMC1 and RAD51 foci were elevated compared to wild-type littermates. Foci numbers increased to maximal levels in early zygotene with a ∼1.5-fold higher than normal average number. At this stage, nearly all mutant cells had higher numbers of DMC1 and RAD51 foci (227–462 and 286–452 foci for DMC1 and RAD51, respectively) compared to wild type (155–315 and 179–322 foci for DMC1 and RAD51, respectively). Foci numbers declined in mid zygotene, as in wild type, but most mutant cells continued to harbour elevated numbers of foci compared to wild type. Foci further declined in late zygotene-like and pachytene-like cells, but not to same extent as in wild type. Importantly, DMC1 and RAD51 foci were prevalent on synapsed axes in mutant pachytene-like cells, consistent with the retention of γH2AX, while synapsed wild-type pachytene chromosomes had substantially fewer foci (**Figure 5B, Supplementary Figure 3A**). We noticed that individual DMC1 foci appeared to be visibly brighter in *Mcm8^m/m^* compared wild type (**Figure 5B**); this was also reported for *Mcm8^-/-^* mice (Lutzmann et al. 2012). Possible explanations for this are described in the discussion section.

RPA foci generally showed a similar trend as DMC1/RAD51 foci, with a few important differences (**Figure 5E, Supplementary Figure 3B**). In wild type, RPA foci accumulate during leptotene and early zygotene reaching highest levels in mid zygotene and decline over later stages. RPA foci numbers were elevated on average in *Mcm8^m/m^* leptotene and early zygotene compared to wild type, but showed a wide range, such that many mutant cells were similar to wild type. In mid zygotene, RPA foci numbers were similar or slightly decreased in mutants compared to wild type, although this difference was not statistically significant. Foci remained elevated in *Mcm8^m/m^* pachytene-like cells, while they declined to lower numbers in wild type.

To summarize, *Mcm8^m/m^* spermatocytes accumulate higher numbers of DMC1 and RAD51 foci during leptotene and early zygotene, suggesting that mutant spermatocytes form higher levels of DSB-associated ssDNA. Consistent with this interpretation, RPA foci are also elevated in leptotene and early zygotene, albeit to a lesser extent. We attribute the more modest elevation in RPA foci, along with the reduction in RPA foci numbers in mutant mid zygotene to a decrease in recombination intermediate-associated ssDNA (see below). Recombination foci generally decline over later stages, indicating that some repair ensues. New DSBs may also form on persistently asynapsed regions (Kauppi et al. 2013), contributing to foci counts. However, *Mcm8^m/m^* spermatocytes that achieve substantial levels of autosomal synapsis (pachytene-like) display flares of γH2AX, as well as elevated numbers of DMC1, RAD51 and RPA foci on synapsed axes. We interpret these as signs of incompletely repaired DSBs. We conclude that meiotic recombination is substantially dysregulated in MCM8-deficient spermatocytes. Furthermore, the recombination defects along with the synaptic defects we describe here likely account for the spermatogenic arrest during prophase in MCM8-deficient mice.

### *Mcm8^m/m^* spermatocytes form more SPO11-dependent DSBs

We hypothesized that the elevated recombination foci in *Mcm8^m/m^*leptotene and early zygotene are due to increased DSBs. We therefore quantified γH2AX during meiotic prophase. The increase in DSBs may be caused by increased formation of meiotic breaks or may reflect meiotic replication-associated damage that is carried over into prophase, so we also quantified γH2AX in preleptotene cells that are undergoing meiotic replication.

γH2AX staining was barely detectable in preleptotene cells (**Figure 6A**) and the intensity of immunofluorescence signal was comparable between wild-type and *Mcm8^m/m^* preleptotene cells (**Figure 6B**). In wild-type prophase, γH2AX signal intensity increased in leptotene through early zygotene concurrent with break formation, and then declined in mid zygotene (**Figure 6A, B**), as expected (Mahadevaiah et al. 2001). γH2AX intensity did not diminish further in later stages in wild type, partly due to the appearance of DSB-independent γH2AX on sex chromosomes. *Mcm8^m/m^* spermatocytes also acquired γH2AX in leptotene and early zygotene, but more than wild type (**Figure 6A**). Average signal intensities were 1.9-fold and 1.4-fold higher than normal in leptotene and early zygotene, respectively (**Figure 6B**). γH2AX signal declined on average in mid zygotene but remained higher than wild type, including in late zygotene-like and pachytene-like cells, consistent with a DSB repair defect.

**Figure 6.**
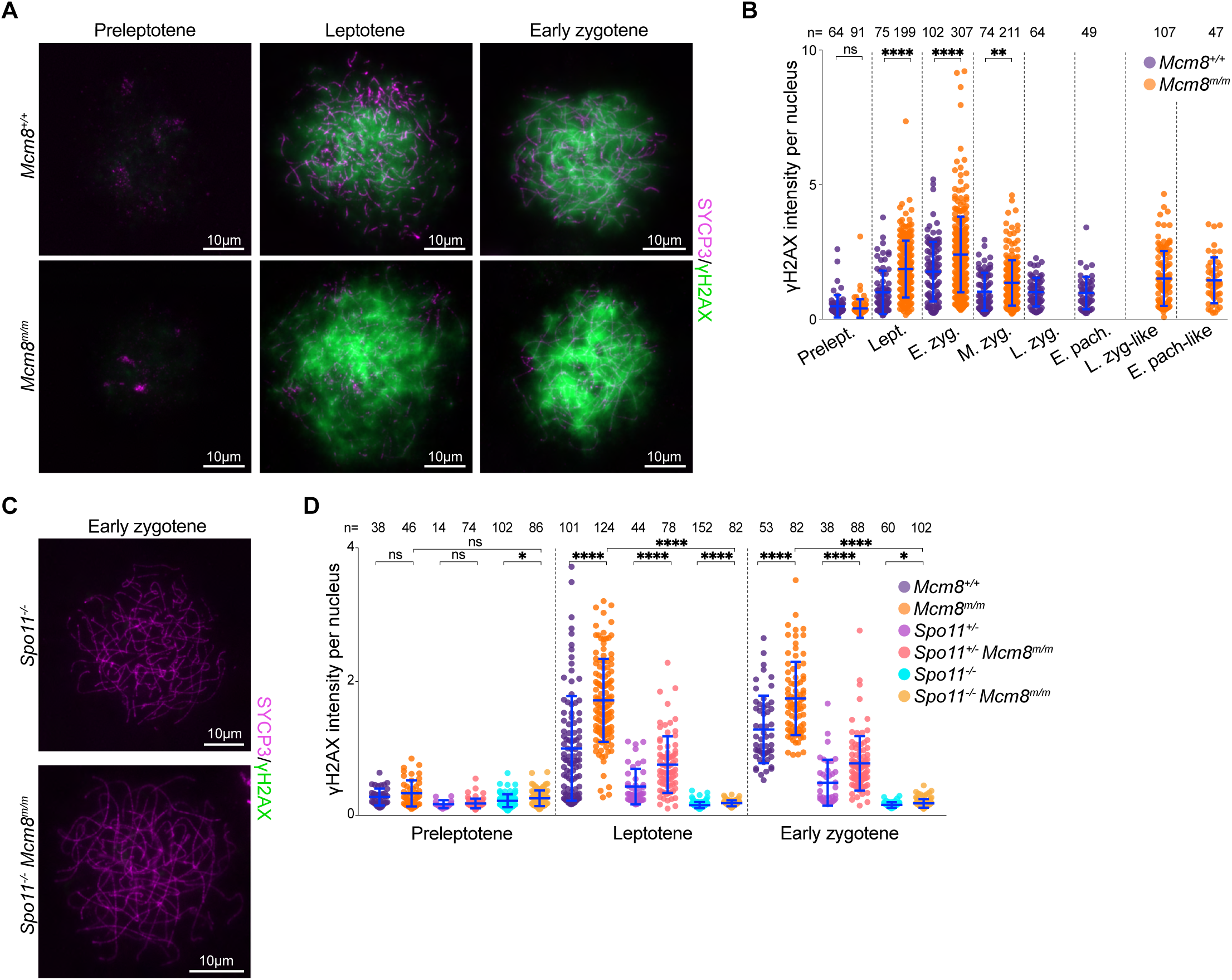
MCM8 deficiency leads to increased SPO11-dependent DSBs. **(A)** Spermatocyte spreads immunostained for SYCP3 and γH2AX. (**B**) Quantification of γH2AX intensity. (**C**) Chromosome spreads of early zygotene cells immunostained for SYCP3 and γH2AX. (**D**) γH2AX intensity measurements during preleptotene, leptotene and early zygotene. In (**B**) and (**D**), each point represents the signal intensity measurement for one cell, normalized to the mean intensity of wild-type leptotene. Means ± standard deviations for three mice are depicted using blue lines. Results of two-tailed Mann Whitney tests are shown: ns is non-significant (p > 0.05), * is p ≤ 0.05; ** is p ≤ 0.01, and **** is p ≤ 0.0001. Meiotic stages were assessed based on SYCP3-staining patterns: preleptotene (Prelept.), leptotene (Lept.), early zygotene (E. Zyg.), mid zygotene (M. Zyg.), late zygotene (L. Zyg.), early pachytene (E. Pach.), late zygotene-like (L. Zyg-like), early pachytene-like (E. Pach-like).

We infer from the lack of visible γH2AX during preleptotene and the increase in γH2AX during leptotene/early zygotene that MCM8 deficiency leads to increased levels of meiotic DSBs. If so, we expect the γH2AX signal to be dependent on the presence of SPO11, which catalyzes meiotic DSBs during leptotene and early zygotene (Baudat et al. 2000; Romanienko and Camerini-Otero 2000; Lam and Keeney 2014). Indeed, *Mcm8^m/m^* spermatocytes that also lacked SPO11 (*Spo11^-/-^Mcm8^m/m^*) lacked visible γH2AX staining (**Figure 6C**). While the mean γH2AX signal intensities were elevated in *Spo11^-/-^Mcm8^m/m^* compared to *Spo11^-/-^* alone (**Figure 6D, Supplementary Figure 4**), possibly indicating the presence of some SPO11-independent DSBs, the increase was extremely marginal (1.17-, 1.19- and 1.15-fold average increase in preleptotene, leptotene and early zygotene, respectively). *Spo11^+/-^*heterozygotes form fewer DSBs and therefore accumulate lower levels of γH2AX compared to wild type (Lange et al. 2011; Cole et al. 2012). MCM8 deficiency in these cells (*Spo11^+/-^Mcm8^m/m^*) led to an elevation of γH2AX levels (**Figure 6D**), in keeping with the elevation seen in *Mcm8^m/m^* where *Spo11* is wild type. Taken together, these results suggest that MCM8 deficiency leads to an increase in SPO11-dependent DSBs.

### MCM8 depletion affects the formation and/or stability of recombination intermediates downstream of DSB resection

Because *Mcm8^m/m^* spermatocytes form extra meiotic breaks and fail to repair these breaks, we sought to determine whether breaks form at expected locations and whether the initial steps of recombination occur normally. SPO11-induced DSBs are resected to generate ssDNA, which are bound initially by RPA and subsequently by the recombinases DMC1 and RAD51, creating a nucleoprotein filament that engages in homology search (Brown and Bishop 2014; Raina et al. 2025). Subsequent strand invasion generates a displaced loop of DNA (D-loop intermediate), which can be channeled through different pathways that result in DSB repair (Hunter 2015). We examined each of these events in detail.

We first mapped break sites genome-wide by sequencing DMC1-bound ssDNA (ssDNA sequencing or SSDS (Brick et al. 2018)) from *Mcm8^m/m^* and *Mcm8^+/+^*adult testes. SPO11-induced DSBs occur primarily at hotspots whose locations have been previously mapped in wild-type B6 mice (Brick et al. 2012; Lange et al. 2016). Most DSB hotspots identified by DMC1-SSDS in *Mcm8^m/m^* and *Mcm8^+/+^* overlapped with previously mapped sites (**Figure 7A**). We think that the non-overlapping hotspots likely reflect the mixed B6/FVB background of our mice and are due to documented strain-specific differences in DSB locations (Smagulova et al. 2016). Importantly, hotspots were largely shared between *Mcm8^m/m^* and *Mcm8^+/+^* littermates (93% and 75% of hotspots in *Mcm8^+/+^* and *Mcm8^m/m^*, respectively; **Figure 7A**). Moreover, the relative usage of individual hotspots, inferred by the strength of DMC1 SSDS signal within peaks, was highly correlated in *Mcm8^m/m^*and *Mcm8^+/+^* at autosomes (**Supplementary Figure 5**). Hotspots on sex chromosomes were less enriched in *Mcm8^m/m^* compared to controls. DSBs formed on nonhomologous regions of the sex chromosomes are thought to persist longer due to delayed repair (Lange et al. 2016), also evidenced by persistent DMC1 foci on these regions during pachytene (Moens et al. 2002), so the lower DMC1-SSDS signal for sex chromosomes in *Mcm8^m/m^*is likely explained by the depletion of pachytene cells in adult mutant testes. We conclude that MCM8 deficiency does not substantially alter meiotic DSB locations.

**Figure 7.**
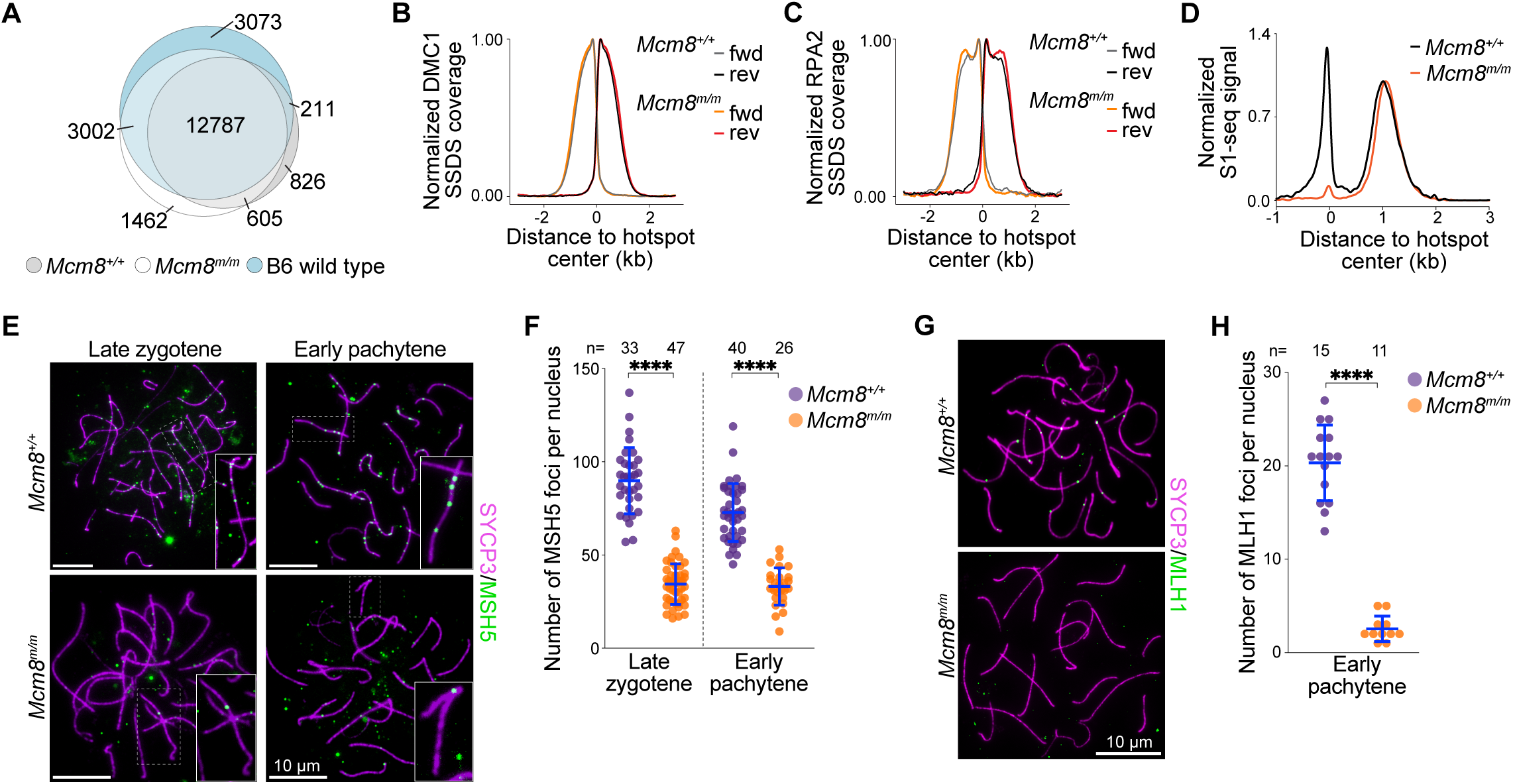
MCM8 functions downstream of resection to regulate recombination intermediates. **(A)** Overlap of hotspots identified by DMC1 SSDS in adult mice of the indicated genotypes. B6 wild type refers to B6 strain-specific hotspots. (**B**) Metaplots of DMC1 SSDS averages around B6 wild-type hotspots in adult mice. Sequencing reads originating from forward (fwd) and reverse (rev) strands are shown separately. Strand-specific reads were averaged and normalized by setting the highest value of the resection peak to 1. (**C**) Metaplots of RPA2 SSDS averages around B6 wild-type hotspots in adult mice. Reads originating from forward (fwd) and reverse (rev) strands are shown separately. (**D**) Genome-average S1-seq at B6 wild-type hotspots in adult mice. Reads from the reverse strand were flipped and combined with forward strand reads, averaged, and normalized by setting the height of the resection peak to 1. (**E**) Chromosome spreads immunostained for SYCP3 and MSH5. Insets show higher-magnification views of foci on chromosome axes. (**F**) Quantification of MSH5 foci in late zygotene and early pachytene-like cells. (**G**) Chromosome spreads of early to mid pachytene-like cells immunostained for SYCP3 and MLH1. (**H**) Quantification of MLH1 foci in pachytene-like cells. In (**F**) and (**H**), foci overlapping with SYCP3-positive axes were scored. Each point is a count from an individual cell. Means ± standard deviations for three mice are depicted in blue lines. Results of two-tailed Mann Whitney tests are shown: ****, p ≤ 0.0001. Staging was done using SYCP3-staining patterns, where early pachytene stage cells were defined as those with fully synapsed autosomes along with extensive X-Y synapsis.

The spatial distribution of ssDNA binding proteins around DSB hotspots provides information about their localization as well as the length of ssDNA generated (Hinch et al. 2020). We therefore examined DMC1 binding profiles at hotspots. Distributions of DMC1-SSDS signal around hotspots in *Mcm8^m/m^* and *Mcm8^+/+^*were closely aligned (**Figure 7B**) and were consistent with previous descriptions of DMC1-SSDS in wild-type mice (Brick et al. 2012). DMC1-SSDS signal was maximal adjacent to hotspots and extended to ∼1.8 kb away from hotspots, with strand polarity consistent with binding to 3’ ssDNA overhangs on either side of DSBs (Khil et al. 2012).

We also examined RPA binding at hotspots by sequencing RPA-bound ssDNA (RPA-SSDS (Hinch et al. 2020)) from adult testes. RPA-SSDS profiles were also mostly overlapping in *Mcm8^m/m^*and *Mcm8^+/+^* (**Figure 7C**). RPA-SSDS signal extended to ∼2 kb on either side of DSBs with strand polarity that is consistent with binding 3’ ssDNA overhangs in both genotypes (Hinch et al. 2020). The wider footprint of RPA signal compared to DMC1 signal is expected and is due to differences in localisation of ssDNA binding proteins: DMC1 binds ssDNA closer to DSB ends, RAD51 binds more distally, and RPA binds across the length of the ssDNA. RPA signal was highest close to hotspots in both *Mcm8^m/m^* and *Mcm8^+/+^*, but more distally the signal accumulated to higher levels forming a more prominent shoulder in *Mcm8^m/m^*. This may reflect a biologically meaningful difference in RPA accumulation along ssDNA or an experimental artifact due to better signal-to-noise ratio in mutant testes that harbour elevated numbers of incompletely repaired DSBs (see below).

RPA binds ssDNA at nascent 3’ overhangs created by resection as well as ssDNA within recombination intermediates, so RPA-SSDS captures RPA localization at both ssDNA substrates (Hinch et al. 2020). In wild type, strand-specific RPA signal associated with 3’ ssDNA on one side of the DSB extends into the opposite side of the DSB. This signal extension is thought to come at least in part from RPA binding to ssDNA on the repair template, ie, RPA binding to DSB-proximal ssDNA in a recombination intermediate such as a D-loop, possibly after some amount of extension and/or further unwinding of the template DNA duplex. Interestingly, this pattern of DSB-proximal RPA-SSDS reads that have the wrong polarity to be resection tracts was present in wild type, but it was not apparent in *Mcm8^m/m^* (**Supplementary Figure 5B**).

Our SSDS experiments suggest that population-average lengths of DMC1-coated and RPA-coated ssDNA at resected DSBs does not appear to be substantially altered in MCM8-deficient mice. The SSDS experiments also suggest that RPA-coated ssDNA at recombination intermediates may be depleted in MCM8-deficient mice. We reasoned that the length of ssDNA available is likely to be normal in MCM8-deficient mice. We further reasoned that fewer recombination intermediates may be present in mutant mice. To test these hypotheses, we performed S1-sequencing (S1-seq) (Yamada et al. 2020; Kim et al. 2024) in *Mcm8^m/m^*and *Mcm8^+/+^* adult testes. In this method genomic DNA is digested with ssDNA-specific nuclease S1 and the blunted DNA ends are sequenced. This provides a population-averaged measurement of genome-wide resection endpoints, irrespective of the proteins bound there. Additionally, S1-seq generates reads adjascent to DSB hotspot centres that are thought to arise from recombination intermediates, inferred to be D-loops (Yamada et al. 2020).

S1-seq yielded sequencing reads distributed ∼0.3–2 kb from DSB hotspots with polarity that is consistent with resection tracts (**Figure 7D**) (Yamada et al. 2020). Resection-associated S1-seq profiles were similar in *Mcm8^m/m^*and *Mcm8^+/+^*, although mutants showed a modest increase in the proportion of sequencing signal from longer tracts (**Supplementary Figure 5C**). Resection tracts in wild type averaged 1075 bp, consistent with previous findings (Yamada et al. 2020). Resection lengths in *Mcm8^m/m^*averaged 1144 bp (6.4% longer than wild-type average), suggesting that a subset of DSBs in *Mcm8^m/m^* may be resected more. It’s possible that MCM8 limits the extent of resection. We consider it more likely that this modest increase in *Mcm8^m/m^* reflects the accumulation of cells that are unable to proceed with recombination (see below) and/or cell population differences between adult mutant and wild-type testes.

Strikingly, the S1-seq signal corresponding to recombination intermediates was largely absent in *Mcm8^m/m^* mice (**Figure 7D, Supplementary Figure 5D**). This is consistent with the diminished recombination intermediate-associated RPA foci in mid zygotene and suggests that *Mcm8^m/m^* harbour fewer recombination intermediates. The loss of this DSB-proximal S1-seq signal has previously also been observed in *Dmc1^-/-^* mice that are unable to perform strand exchange and therefore lack intermolecular recombination intermediates (Brown and Bishop 2014; Yamada et al. 2020). MCM8-deficient cells may form fewer recombination intermediates, or recombination intermediates may be unstable, or intermediates may turn over more rapidly. We also cannot exclude the possibility that intermediates form but adopt a configuration that makes them insensitive to S1 nuclease or makes them difficult to detect in our assay.

Recombination intermediates such as D-loops are bound and stabilized by meiotic ZMM proteins, including the MSH4-MSH5 heterodimer, a subset of which are processed into crossovers by the MLH1-MLH3 complex (Gray and Cohen 2016; Pyatnitskaya et al. 2019; Frasca et al. 2025). We surmised that the depletion of recombination intermediates in *Mcm8^m/m^*mice may be accompanied by a decrease in MSH4-MSH5-bound recombination sites, and consequently also lead to a decrease in MLH1-MLH3-bound sites. We therefore examined the behaviour of these recombination proteins.

In wild type, axes-associated MSH5 foci were abundant in late zygotene and foci numbers declined in early pachytene (**Figure 7E**, **F**), consistent with previous reports analysing MSH4-MSH5 foci (Kneitz et al. 2000; Moens et al. 2002; Moens et al. 2007). MSH5 foci numbers were strongly reduced in *Mcm8^m/m^* spermatocytes at both stages examined (∼2.6-fold and ∼2.2-fold lower than normal averages in late zygotene-like and early pachytene-like, respectively). Axes-associated MLH1 foci were more severely depleted (**Figure 7G**, **H**). *Mcm8^m/m^*early pachytene-like spermatocytes with apparently normal autosomal synapsis harboured only very few MLH1 foci (1–5 foci), while higher numbers of foci (average 20 foci) were present in wild-type early/mid pachytene cells, as expected (Plug et al. 1998; Anderson et al. 1999; Moens et al. 2002; Moens et al. 2007).

To summarize, SPO11-dependent DSBs form at hotspots and are mostly resected normally in *Mcm8^m/m^* spermatocytes. Resected 3’ ssDNA accumulate DMC1. Downstream recombination intermediates are barely detected, however. Recombination intermediate-associated RPA is also depleted, both in our population-averaged SSDS assay and in our cytological assay. MSH4-5-marked stable recombination sites are severely reduced, and therefore, not surprisingly, MLH1-bound late recombination sites are nearly absent. We posit that MCM8 is required for the formation and/or stability of post-resection recombination intermediates during meiosis.

### MCM8 binds D-loop structures

We next sought to examine the DNA binding preferences of MCM8. Previous biochemical studies have primarily examined the DNA binding specificity of the MCM8-9 complex prepared by co-expression and co-purification of the two proteins (Huang et al. 2020; Acharya et al. 2024). Under these conditions, MCM8 and MCM9 assemble into a heterohexamer with DNA unwinding activity. Individually expressed proteins are largely monomeric and do not show DNA unwinding activity (Acharya et al. 2024). Because the meiotic phenotypes of *Mcm8^-/-^* and *Mcm9^-/-^* mice are only partially overlapping (Hartford et al. 2011; Lutzmann et al. 2012), MCM8 may not necessarily function exclusively in a complex with MCM9 during meiotic recombination. We therefore examined the DNA binding preferences of MCM8-9 as well as MCM8 alone. Additionally, our *in vivo* data suggest that MCM8 may function on post-resection recombination intermediates. Therefore, we examined MCM8-9 and MCM8 binding to different types of oligonucleotide-based D-loop substrates.

The MCM8-9 hexamer has been reported to preferentially bind and unwind branched DNA structures (Acharya et al. 2024). Overhanging DNA is bound by MCM8-9 slightly less efficiently, and fully double-stranded DNA substrates and single-stranded DNA are the least preferred (Acharya et al. 2024). We could recapitulate these findings (**Figure 8**, **Supplementary Figure 6**), and additionally show that D-loop structures are also excellent substrates for MCM8-9. MCM8 alone bound DNA overall less efficiently than the MCM8-9 complex and displayed an altered DNA substrate preference (**Figure 8**, **Supplementary Figure 6**). MCM8 clearly preferred D-loop structures relative to the other DNA substrates tested. Our analysis of the D-loop structure binding suggests that MCM8 prefers a flap at the junction point between double-stranded and single-stranded DNA. In contrast to MCM8-9, the 3′ overhang does not appear to be a suitable substrate for MCM8 alone, as binding reaches only about 20% even at the highest protein concentration of 150 nM (**Figure 8**, **Supplementary Figure 6**). Overall, these results support the hypothesis that MCM8 may preferentially associate with D-loop intermediates formed after DNA end resection.

**Figure 8.**
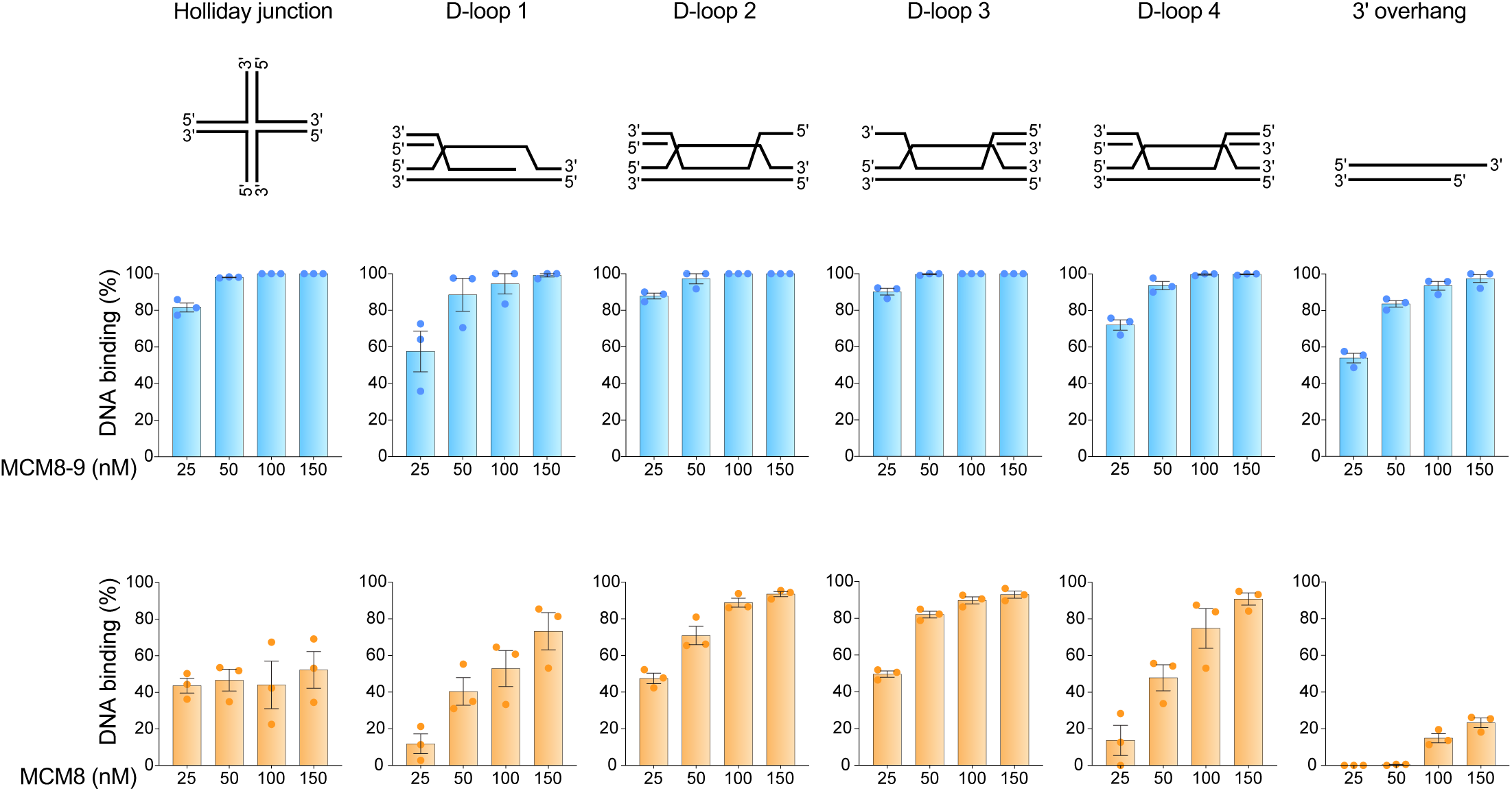
DNA binding preferences of MCM8 compared to MCM8-9. DNA-binding assays using the indicated substrates. Top: schematics of substrates used. Bottom: quantitation of DNA binding (electrophoretic mobility shift assays shown in **Supplementary** Figure 6) for MCM8-9 in blue and MCM8 alone in yellow. Bars show averages and standard error of the mean for three independent experiments.

## Discussion

MCM8 was previously shown to be essential for meiotic recombination and therefore gametogenesis in mice (Lutzmann et al. 2012), but mechanistic detail was lacking. Moreover, it was unclear to what extent the role of MCM8 in mammalian meiotic recombination is analogous to that proposed for its fly ortholog, REC. Here we implicate MCM8 in controlling mammalian meiotic recombination in at least two ways. MCM8 limits meiotic DSB numbers and regulates recombination intermediates. Possible scenarios of how MCM8 may accomplish these roles and comparisons with its proposed role in flies are discussed below.

### Role in regulating meiotic DSBs

How MCM8 controls DSB frequency is unclear. DSBs activate the ataxia-telangiectasia mutated (ATM) protein kinase, which phosphorylates H2AX and suppresses additional DSB formation via a negative feedback loop (Lange et al. 2011; Keeney et al. 2014; Lukaszewicz et al. 2018), but it is unlikely that MCM8 contributes to ATM-dependent inhibition of DSBs. Perturbation of ATM signaling typically results in clusters of DSBs that are not resolvable as cytologically distinct recombination foci (Lange et al. 2011; Garcia et al. 2015; Lange et al. 2016), while *Mcm8^m/m^* spermatocytes form more RAD51 and DMC1 foci. Additionally, ATM signaling loss leads to increased breaks in normally weak hotspots (Lange et al. 2016; Yamada et al. 2017), but we detect no obvious alterations in hotspot usage in *Mcm8^m/m^*mutants. Our findings are also inconsistent with a model in which an asynapsis-triggered positive feedback loop leads to continued DSB formation in mutants (Kauppi et al. 2013), as these additional breaks would be expected to occur during late prophase while *Mcm8^m/m^* harbor extra DSBs early in prophase.

One possibility is that MCM8 affects DSB frequency through an activity during meiotic replication, which is known to be spatiotemporally coupled with the subsequent formation of meiotic DSBs (Borde et al. 2000; Murakami and Keeney 2014; Pratto et al. 2021). Consistent with this hypothesis, we detect MCM8 in meiotic replicating cells, although MCM8 appears to be non-essential for completing meiotic replication. During or shortly after meiotic replication, MCM8 may inhibit localization of pro-DSB factors or may influence meiotic chromosome organization, which in turn could impact meiotic DSB numbers (Kumar et al. 2010; Stanzione et al. 2016; Tesse et al. 2017; Kumar et al. 2018; Boekhout et al. 2019; Papanikos et al. 2019; Grey and de Massy 2021; Dereli et al. 2024).

### Role in meiotic DSB repair

After resection and strand exchange, D-loops are bound and stabilized by MSH4-5, prior to all interhomolog repair (Snowden et al. 2004; Pyatnitskaya et al. 2019; Frasca et al. 2025). MCM8 mutants fail to accumulate normal levels of MSH4-5, and *Mcm8* mutant phenotypes resemble those in *Msh4^-/-^* and *Msh5^-/-^*, namely accumulation of DMC1 and RAD51 foci, severely compromised synapsis, impaired DSB repair, and spermatocyte death during mid-prophase (de Vries et al. 1999; Edelmann et al. 1999; Kneitz et al. 2000). Thus, MCM8 may act upstream of or alongside MSH4-5 to stabilize recombination intermediates. At D-loops, MCM8 may be important for DNA synthesis of the invading strand, akin to its proposed function in cell lines (Natsume et al. 2017; Hustedt et al. 2019; Huang et al. 2020), without which D-loops may be more transient. We note that the recombination intermediate-associated S1-sequencing signal that is lost in *Mcm8^m/m^* is thought to represent D-loops where the 3’ end of the ssDNA is not fully invaded, possibly even capped by SPO11-oligos complexes, and therefore is not available to prime DNA synthesis (Yamada et al. 2020). If so, absence of this signal in mutants suggests that MCM8 acts on strand-exchange intermediates prior to DNA synthesis, and a role in extension of the invading strand/D-loop would therefore be a secondary function of MCM8. Our *in vitro* data show that MCM8 preferentially binds D-loop structures and is consistent with these possibilities.

Curiously, while resection tract lengths appear mostly normal and DMC1 accumulation on ssDNA as assessed by DMC1-SSDS is also normal, individual DMC1 foci are brighter across prophase in *Mcm8^m/m^* than in wild type (Lutzmann et al. 2012). Individual RAD51 foci are also brighter in mutants, although to a lesser extent than DMC1. This phenotype partially resembles that described for *Fignl1* and *Firrm* mutants where impaired regulation of DMC1/RAD51 accumulation appears as altered, sometimes extended DMC1/RAD51 staining patterns at recombination sites when visualized by super resolution microscopy, and precludes formation of stable MSH4-5-bound recombination intermediates (Ito et al. 2023; Zhang et al. 2023; Zainu et al. 2024). Alterations in the architecture of DMC1/RAD51 nucleofilaments upon loss of MCM8-mediated stabilization of D-loops may make DMC1/RAD51 foci appear brighter by conventional microscopy. However, we cannot exclude the possibility that MCM8 plays a role in regulating DMC1/RAD51 nucleofilaments in a manner that does not alter population-averaged measurements of DMC1 occupancy and resection. In this scenario, MCM8 may directly or indirectly facilitate homology search or strand invasion to form the D-loop. Yet another non-exclusive possibility is that MCM8 facilitates stable D-loop formation by disassociating inappropriate recombination intermediates, such as those formed between non-allelic sequences.

Synaptic defects in *Mcm8^m/m^* are likely attributable to impaired recombination and resemble those seen in mutants that are defective for forming or stabilizing recombination intermediates, such as *Dmc1^-/-^*, *Hop2^-/-^*, *Msh4^-/-^* and *Msh5^-/-^* (Pittman et al. 1998; Yoshida et al. 1998; de Vries et al. 1999; Edelmann et al. 1999; Kneitz et al. 2000; Petukhova et al. 2003; Pezza et al. 2006; Yamada et al. 2020). Moreover, many *Mcm8* mutant spermatocytes form short stretches of the synaptonemal complex, and few cells even achieve complete autosomal synapsis, indicating that MCM8 is not required for synaptonemal complex formation *per se*. *Mcm8^m/m^* spermatocytes with synaptic defects and unrepaired DSBs are likely eliminated by the recombination checkpoint and/or due to sex body failure (Royo et al. 2010; Pacheco et al. 2015). Those with apparently normal autosomal synapsis nonetheless harbour unrepaired DSBs, therefore would suffer the same fate.

### MCM8-containing complexes

It is unlikely that MCM8 functions exclusively as part of a heterohexameric helicase with MCM9 in mouse meiotic cells, as individual loss of these proteins has only partially overlapping meiotic phenotypes. While MCM9 loss has been reported to disrupt meiotic recombination, some *Mcm9^-/-^*spermatocytes nonetheless form sperm and *Mcm9^-/-^* males are fertile (Hartford et al. 2011; Lutzmann et al. 2012). One explanation could be that MCM8 functions both as part of the MCM8-MCM9 helicase complex as well as independent of MCM9, and the meiotic defects we report here are a result of losing both activities. The specific meiotic recombination defects in *Mcm9^-/-^* spermatocytes are unexplored and may differ from those in *Mcm8* mutants. Therefore, we cannot exclude the possibility that MCM8 and MCM9 function independently in meiotic cells.

The *Mcm8* mutant phenotypes we report here resemble those seen in *Mcmdc2^-/-^* mice, in that both mutants have severely compromised synapsis, and both accumulate DMC1/RAD51 foci but MSH4-5-marked recombination sites are diminished leading to male infertility (Finsterbusch et al. 2016; McNairn et al. 2017). MCMDC2 resembles typical MCM proteins in domain architecture and is the mammalian ortholog of *Drosophila* MEI-217 and MEI-218 (Kohl et al. 2012; Finsterbusch et al. 2016; McNairn et al. 2017). These fly proteins encode the N-terminal and C-terminal domains of MCMDC2 and physically interact with each other, as well as with the fly ortholog of MCM8. Thus, MCM8 may function in conjunction with and possibly in a complex with MCMDC2 during mouse meiosis, like in flies.

Meiotic functions of MCM8 that are independent of MCM9 may not involve DNA unwinding. Recombinant MCM8 is largely monomeric and does not exhibit DNA unwinding activity *in vitro* when expressed without MCM9 (Acharya et al. 2024). Additionally, MCMDC2 harbors amino acid changes within its ATPase domain that are predicted to disrupt ATPase activity (Kohl et al. 2012), thus a hypothetical MCM8-MCMDC2 complex may exhibit reduced unwinding activity or no unwinding activity at all. It is also possible that MCM8 interacts with other MCM proteins to form a functional helicase during meiosis. In flies, a meiosis-specific mutation in the replicative helicase subunit MCM5 leads to a meiotic recombination defect similar to that observed upon loss of the MCM8 ortholog, leading to the proposal that these MCMs may form a complex during fly meiosis (Lake et al. 2007; Kohl et al. 2012). Whether replicative MCMs serve a role during mammalian meiotic recombination is currently unknown.

### Roles of MCM8 in flies vs mice

The meiotic role of MCM8 was first described in flies, where a complex called Mei-MCM composed of MCM8 (fly REC) and MCMDC2 (fly MEI-217 and MEI-218) was proposed to antagonize activities that dismantle recombination intermediates (Blanton et al. 2005; Kohl et al. 2012). It was further proposed to functionally replace MSH4-5, which is important for stabilizing recombination intermediates in mice, but is absent in flies. Our findings suggest that like flies, mice also require MCM8 to form stable recombination intermediates.

The consequence of losing this activity differs in flies and mice, however. In mice, loss of MCM8 results in a failure to repair DSBs, as does the loss of MSH4-5 (de Vries et al. 1999; Edelmann et al. 1999; Kneitz et al. 2000; Milano et al. 2019; Frasca et al. 2025). Repair failure is accompanied by severe synapsis defects. MCMDC2 loss also has similar consequences (Finsterbusch et al. 2016; McNairn et al. 2017). In flies, loss of Mei-MCMs leads to reduced crossovers but DSBs are repaired as noncrossovers (Blanton et al. 2005). This may be because interhomolog pairing and synapsis are independent of meiotic DSBs in flies, and the synaptonemal complex may sufficiently stabilize recombination intermediates for noncrossover repair events, as previously proposed (Lake and Hawley 2012; McNairn et al. 2017). While in mice, where meiotic DSBs mediate pairing and synapsis, both noncrossover and crossover repair may require the D-loop stabilizing activities of MCM8 and MSH4-5 (McNairn et al. 2017; Zickler and Kleckner 2023; Frasca et al. 2025).

REC/MCM8 containing complexes may also be similar in flies and mice. Flies lack MCM9 and mouse MCM8 partly functions independently of MCM9 during meiosis, at least for functions that are essential to meiotic recombination (Blanton et al. 2005; Hartford et al. 2011; Lutzmann et al. 2012). Additionally, phenotypic similarities between *Mcm8* mutant and *Mcmdc2* mutants suggest that these proteins collaborate functionally, perhaps even associate physically, similar to REC, MEI-217 and MEI-218 in flies (Kohl et al. 2012; Finsterbusch et al. 2016; McNairn et al. 2017).

REC is highly diverged in Schizophoran flies (Blanton et al. 2005; Liu et al. 2009; Kohl et al. 2012). This coupled with the loss of MCM9 led to the idea that MCM8 evolved a novel function in this taxa to facilitate meiotic recombination (Kohl et al. 2012). We propose an alternative hypothesis that MCM8-containing MCM complexes serve a more ancestral role in recombination intermediate processing. Existing activities may have been modified during evolution, in that flies lost MSH4-5 and utilized only MCMs to stabilize meiotic recombination intermediates, while the role of MCMs in mice was retained. Any additional roles in mammals (for example, roles that impact meiotic DSB frequency or roles in mitosis) may also be ancestral or may reflect functional expansion of MCM8-containing complexes. A related hypothesis was proposed for MCMDC2-containing complexes (Finsterbusch et al. 2016).

Although MCM8 appears to have been lost in some taxa, including in some fungi and some animals, it is present in most eukaryotic lineages, including deeply branching eukaryotes (for example, some excavates), suggesting that it likely arose early in eukaryotic evolution (Blanton et al. 2005; Liu et al. 2009; Kohl et al. 2012). On the basis of the functional similarities between flies and mice, and because MCM8 is widely distributed in eukaryotes, we propose that MCM8 is an evolutionarily ancient regulator of recombination-mediated DNA repair and genome stability.

## Materials and Methods

### Mice

Experimental procedures involving mice were performed following regulatory standards and were approved by the Rutgers University Institutional Animal Care and Use Committee. Animals were housed in a room with 12-hour light/dark cycle, 70°–74°F ambient temperature and 30–70% humidity. Mice were fed regular rodent chow and given ad libitum access to food and water. The *Spo11^-^* allele has been described previously (Baudat et al. 2000). *Mcm8* mutant animals harbor a mixed inbred background of C57BL/6J (Jackson Laboratory) and FVB/NJ (Jackson Laboratory).

The *Mcm8^mara^* (*Mcm8^m^*) allele was generated through a random mutagenesis screen designed to isolate autosomal recessive mutations, as previously described (Jain et al. 2017; Jain et al. 2018). Briefly, male C57BL/6J (B6) mice were mutagenized with *N*-ethyl-*N*-nitrosourea (ENU) and subsequently crossed with wild-type FVB/NJ (FVB) female mice, producing founder males that are potential carriers of a mutation of interest. Founder males were bred with wild-type FVB females to produce second-generation daughters, half of which are expected to be mutation carriers. Second-generation daughters were crossed back to founder males, producing third-generation offspring that are potentially homozygous for the mutation. Third generation males aged 16 to 20 dpp were screened for meiotic defects and mouse lines displaying defects were maintained. Screening was performed at Memorial Sloan Kettering Cancer Center, New York.

To identify the *mara* mutation, we used a genetic polymorphism-based positional cloning strategy as described previously (Jain et al. 2017; Jain et al. 2018). Because mutagenesis was conducted on the B6 background and outcrossing was performed with FVB mice, the causative mutation was expected to be linked to B6 variants at polymorphic sites. Accordingly, five third-generation mutant males were analyzed using genome-wide SNP genotyping arrays to identify shared regions of homozygosity for B6 variants. This analysis mapped the causative mutation to a single region on Chromosome 2 (Chr 2). Additional third generation mutant males were analyzed for B6 homozygosity at SNPs within this interval using tailored PCR-based genotyping to narrow the mutation containing interval to a ∼7.5 Mbp region on Chr 2. Finally, we performed whole genome sequencing on one mutant mouse and identified a variant within the mapped interval to be the probable causative mutation. The variant is a T to A mutation within the *Mcm8* gene at Chr 2:132,828,750 (Genome Reference Consortium Mouse Build 38 or mm10). Microarray analysis was done at the Genetic Analysis Facility, The Centre for Applied Genomics, The Hospital for Sick Children, Toronto, Canada. Whole genome sequencing was done at the Integrated Genomics Operation, Memorial Sloan Kettering Cancer Center, New York.

Genotyping of *Mcm8^m^* mice was done by PCR amplification of DNA extracted from toe clips using primers Mcm8F (5’-TGGCAATGTTCTGACAGTTTGG) and Mcm8R (5’-TCACAGAGTAAGTCGGGATGC), followed by sanger sequencing with the amplification primers. Alternatively, genotyping was performed by Transnetyx Automated Gentyping Services (www.transnetyx.com) using quantitative fluorescent PCR. Genotyping of *Spo11^-^* mice was performed by PCR amplification of DNA extracted from toe clips using primers SP1F (5’-CTACCTAGATTCTGGTCTAAGC), PRSF2 (5’-CTGAGCCCAGAAAGCGAAGGAA), and SP16R (5’-ATGTTAGTCGGCACAGCAGTAG) (Baudat et al. 2000).

### EdU treatment

Edu treatment was performed as described previously (Sorkin et al. 2025). Briefly, EdU (Invitrogen) was dissolved in 1X Phosphate-Buffered Saline (PBS) to a final concentration of 4 μg/μl and a single dose of 20 μg/g of body weight was administered intraperitoneally to 14-week-old mice. After 2 hr of incubation, animals were sacrificed, and testes were collected.

### Histology

Histological analyses were performed on adult mice aged 5 to 10 weeks, or juvenile mice aged 13 to 16 dpp (**Supplementary Figure 2)**. Testes isolated from mice were fixed in 4% paraformaldehyde (PFA) or in Bouin’s fixative at 4 °C overnight. Bouin’s fixed testis were washed with deionized water for 1 hr at 4 °C with gentle agitation, followed by two 1 hr washes in 70% ethanol. PFA fixed testes were washed twice in 1X PBS at 4 °C for 30 min each, followed by 30 min in 50% ethanol. Fixed testes were stored in 70% ethanol at 4 °C prior to embedding. Fixed testes were embedded in paraffin, cut into 5-μm-thick sections, and mounted on slides by the Rutgers Research Pathology Services core facility. Sections were deparaffinized with three 5 min washes in xylene and rehydrated with two washes each in 100% ethanol for 3 min, 95% ethanol for 2 min, and 70% ethanol for 2 min, then a single wash in deionized water for 2 min.

Immunofluorescence staining of testes sections were performed as described previously (Sorkin et al. 2025) with a few modifications. Antigen retrieval were performed with sodium citrate buffer (10 mM Sodium Citrate, 0.05% Tween 20, pH 6.0) at 95–100 °C for 20 min, after which slides were allowed to cool at room temperature for 20 min. Slides were blocked for 30 min on a shaker in blocking buffer (0.2% bovine serum albumin (BSA), 0.2% gelatin from cold-water fish skin, 0.05% Tween 20 in 1X PBS), and incubated with primary antibodies at 4 °C overnight in a humidity chamber. A hydrophobic pen (Invitrogen) was used to draw a border around the sample to contain the antibody solution. After primary incubation, slides were washed with blocking solution three times for 5 min each on a shaker, then incubated with secondary antibodies at 37 °C for 1 hr. Slides were washed three times with blocking solution for 5 min each. Slides were mounted with a DAPI-containing mounting medium (Vector Laboratories). A coverslip was added and sealed to the slide with quick dry nail polish. The antibodies used for immunostaining analyses are listed in **Supplementary Table 1**.

Periodic acid Schiff (PAS) staining was performed by the Rutgers Research Pathology Services core facility using PAS Kit (Richard-Allan Scientific, Thermo Fisher Scientific) according to manufacturer’s instructions. Briefly, slides were treated sequentially with 0.5% periodic acid, Schiff reagent, modified Mayer’s hematoxylin, and bluing reagent. Slides were then dehydrated using 100% ethanol, cleared in xylene, and mounted using Permount mounting medium.

TUNEL staining was performed using Click-iT Plus TUNEL Assay Kit (Invitrogen) according to manufacturer’s instructions, with minor modifications. Deparaffinized testes sections were permeabilized by incubation in sodium citrate buffer at 95–100 °C for 5 to 8 mins. After cooling down slides for 20 mins, 100 μl of TdT reaction mixture was added to each slide and slides were incubated at 37 °C for 60 min. Following incubation, slides were rinsed in deionized water, washed with a washing solution (3% BSA, 0.1% Triton X-100 in 1X PBS) for 5 min and rinsed again in 1X PBS. Subsequently, 100 μl of the Click-iT Plus reaction cocktail was added to each slide. We used a 1:10 dilution of kit reaction cocktail to minimize background. Slides were incubated for 30 min at 37 °C in the dark. After incubation, slides were washed with washing solution (3% BSA, 0.1% Triton X-100 in 1X PBS) for 5 min and mounted with a DAPI-containing mounting medium (Vector Laboratories). Coverslips were applied and sealed with quick dry nail polish.

EdU staining was performed according to the manufacturer’s instructions using Click-iT EdU Proliferation Kit (Invitrogen).

### Cytology

Cytological analyses were performed on mice aged 8 to 18 weeks. Spermatocyte squashes were prepared as described (Sorkin et al. 2025). Minced testes tubules were fixed in PFA suspension (2% PFA, 1X PBS, 0.15% Triton X-100) for 1 min in 75 μl per 25 mg testes. After fixation, 15 μl of spermatocyte suspension was placed onto slides and pressed down by a coverslip and left for 1 min. Slides were snap frozen in liquid nitrogen and washed in 1X PBS three times for 3 min with gentle agitation. Slides were stained as described (Sorkin et al. 2025) and mounted with mounting medium containing DAPI.

Spermatocyte spreads were performed as described (Kumar et al. 2024). Briefly, testes were punctured, seminiferous tubules were squeezed out, and immediately fixed in 1% PFA (pH 9.2) on ice for 20 min. Tubules were transferred to a depression slide, subjected to hypotonization in 0.5 M sucrose for 5 min, then chopped finely to make a spermatocyte slurry and incubated at room temperature for 5 min. The spermatocytes were spread onto slides using hypotonic extraction buffer (0.5 M Sucrose, 17 mM Citric Acid, 30 mM Tris-HCl pH 8.0, 5 mM ethylenediaminetetraacetic acid (EDTA), 100 mg/ml phenylmethanesulfonylfluoride (PMSF), 0.5 M dithiothreitol (DTT), pH 8.2-8.4). Spermatocytes spread slides were incubated in a humid chamber for 3 to 4 hr and then allowed to dry slowly overnight with the lid slightly open. Slides were washed with deionized water for 5 min and then with 0.4% Photo-Flo (Kodak) twice for 5 min each, then subjected to immunostaining.

Immunofluorescence staining of spermatocyte spreads was performed as described (Kumar et al. 2024). Briefly, slides were washed with 1X TBST (0.05% Triton X-100 in 1X Tris-Buffered Saline (TBS)) for 5 min, then blocked with antibody dilution buffer (10% Normal Goat Serum, 3% BSA, 0.05% Triton X-100 in 1X TBS) two times for 15 min each. After blocking, slides were stained with primary antibodies by incubation at 4 °C overnight. Stained slides were washed with 1X TBST 3 times for 5 min each, then blocked with antibody dilution buffer for 15 min and incubated with secondary antibodies for 1 hr at 37 °C. After secondary incubation, slides were washed three times with 1X TBST for 5 min each. Slides were mounted with a DAPI-containing mounting medium. The coverslip was sealed to the slide with quick dry nail polish. The antibodies used for immunostaining analyses are listed in **Supplementary Table 1**.

### Image analysis

Immunostained testis sections and spermatocyte nuclei images were captured on a Zeiss Observer Z1 using a 63× oil-immersion or 20× objective. Testis sections stained for detecting MCM8 localization (**Figure 2**) were imaged on a Leica TCS SP8 STED confocal microscope using a 63× oil-immersion. PAS- and TUNEL-stained slides were digitized using EVOS M700 Imaging System (Invitrogen) with a 40× objective.

Quantification of cells and tubule types within sections was performed manually using Zen Blue software (Zeiss). Quantification of foci was performed manually using Zen Blue software (Zeiss). γH2AX signal intensity was quantified using the Contour Spline tool within Zen Blue software (Zeiss). A spline-based border was manually drawn around each spermatocyte nucleus, and the Intensity Mean Value for the fluorophore was recorded. For quantification of TUNEL-positive cells, images were visualized using QuPath software (Bankhead et al. 2017) and scored manually.

### RT-PCR

RNA was isolated from testes of mutant mice and control littermates aged 11 to 14 weeks using RNeasy Plus Mini kit (Qiagen) and cDNA synthesis was performed using the Superscript II Reverse Transcriptase kit (Invitrogen) following the manufacturer’s protocol. qPCR was carried out on a QuantStudio 3 Real-Time PCR System (Applied Biosciences), using SYBR Green Universal Master Mix (Applied Biosciences). Primer sets used for amplification of *Mcm8* are primer set 1: qPCR-F1 (5’-TTCAGAGCTTTCCTCTGCCA), qPCR-R1 (5’-GCAGTGGAGCAAATGACCTC) and primer set 2: qPCR-F2 (5’-GCTGAAGAGAACCTGCTCAA), qPCR-R2 (5’-TACCGCCAAAGAGTGCTAAC). RT-PCR products were resolved on a 2–4% agarose gel and visualized with a ENDURO GDS system (Labnet).

### Immunoblot analysis

Analysis of MCM8 protein levels was performed on adult *Mcm8^m/m^*animals and control littermates aged 11 to 12 weeks as previously described (Sorkin et al. 2025) with some modifications. Testes were lysed in RIPA buffer (50 mM Tris-HCl pH 7.5, 150 mM NaCl, 0.1% sodium dodecyl sulfate (SDS), 0.5% sodium deoxycholate (SDC), 1% NP-40) and one cOmplete Protease Inhibitor Cocktail tablet (Roche) per 10 ml of lysis buffer was added before use. Samples were loaded onto a precast 4–15% Mini Tris-Glycine gels (Bio-Rad) and run at 150 V for 60 min. Then a semidry transfer was performed at 15 V for 25 min onto a nitrocellulose membrane. The membrane was rinsed with water and blocked with blocking solution (5% skim milk, 0.1% Tween 20 in 1X TBS) for 1 hr at room temperature, then incubated with primary antibodies at 4 °C overnight. The membrane was washed three times with TBST (0.1% Tween 20 in 1X TBS) for 10 min each, incubated with secondary antibody in blocking solution for 1 hr at room temperature, and washed again three times with 1X TBST for 10 min each. The chemiluminescence signal was captured with a KwikQuant digital western blot detection system (Kindle Biosciences) and visualized using KwikQuant imager software (Kindle Biosciences). Images were processed using Adobe Photoshop. The antibodies used for western blotting analyses are listed in **Supplementary Table 1**.

### SSDS

DMC1 and RPA SSDS was performed on two 6- to 8-week-old *Mcm8^m/m^* animals and one wild-type littermates as described (Brick et al. 2018). 2 μg of DMC1 antibody (custom) and 2 μg of RPA2 antibody (Abcam ab10359) was used to pull down the nucleoprotein filaments. B6 wild-type hotspots are those described in (Brick et al. 2012). B6 wild-type hotspots present in either wild type or mutant were used to generate SSDS metaplots shown in **Figure 7B**, **C** and **Supplementary Figure 5A**, **B**.

### S1-seq

S1-seq was conducted on two *Mcm8* mutant mice and two wild-type littermates aged 10 weeks. Library preparation, sequencing and bioinformatics analyses were performed as previously described (Kim et al. 2024; Kim et al. 2025). DSB hotspots in **Figure 7D** and **Supplementary Figure 5C**, **D** are hotspots previously identified by SPO11-oligo sequencing (Lange et al. 2016).

### Protein purification

Human MCM8 and the MCM8-9 complex were expressed in *Spodoptera frugiperda* 9 (*Sf9*) insect cells using the Bac-to-Bac expression system (Invitrogen) according to the manufacturer’s recommendations. The MCM8 and MCM9 sequences were codon-optimized for expression in *Sf9* cells (Gen9) and cloned into the final expression constructs pFastBac1-MCM9co and pFastBac1-FLAG-MCM8co (Huang et al. 2020; Acharya et al. 2024).

For purification of the MCM8-9 complex, cells were co-infected with both recombinant baculoviruses at a 1:1 ratio. Cell pellets were resuspended in three pellet volumes of lysis buffer containing 50 mM Tris-HCl (pH 7.5), 0.5 mM β-mercaptoethanol, 1 mM EDTA, 1:200 protease inhibitor cocktail (Sigma), 1 mM phenylmethylsulfonyl fluoride (PMSF), 60 µg/ml leupeptin (Merck Millipore), 0.1% NP-40, and cOmplete EDTA-free protease inhibitor cocktail tablets (Roche; 1 tablet per 100 ml). The suspension was incubated for 20 min at 4 °C with continuous stirring. Glycerol was then added to a final concentration of 25%, followed by slow addition of 5 M NaCl to reach a final concentration of 305 mM. The cell suspension was further incubated for 20 min with continuous stirring and subsequently centrifuged at 50,000 ×g for 15 min to obtain the soluble extract. Pre-equilibrated anti-FLAG M2 affinity resin (Sigma) was added to the soluble extract and incubated for 30 min with continuous mixing. The FLAG resin was separated and washed twice in batch with FLAG wash buffer (50 mM Tris-HCl (pH 7.5), 300 mM NaCl, 10% glycerol, 1 mM PMSF, 0.5 mM β-mercaptoethanol, 0.1% NP-40). The resin was then transferred to a disposable column (Thermo Fisher Scientific) and washed for 30 min with the same buffer, followed by an additional wash using FLAG wash buffer containing 100 mM NaCl and no NP-40. Finally, the recombinant protein was eluted from the FLAG resin using FLAG wash buffer (100 mM NaCl) supplemented with FLAG peptide (200 ng/µl; Sigma) and stored at −80 °C. MCM8 was purified using the same procedure.

### DNA substrate preparation

The oligonucleotide-based DNA substrates for the binding assays were prepared by annealing the following oligonucleotides: 3’ overhang (labelled PC1255 and 314), Holliday Junction (labelled PC1253 and PC1254, PC1255 and PC1256), D-loop 1 (labelled BB, BT, InvA, InvB), D-loop 2 (labelled BB, BT, InvA, InvD), D-loop 3 (labelled BB, BT, InvD, InvE), D-loop 4 (labelled BB, BT, InvA, InvD, InvE). The oligonucleotide were 3′-end labelled using T4 Terminal Transferase (New England Biolabs) and [α-^32^P]dCTP (Hartmann Analytic) according to the manufacturer’s instructions and purified with Micro Bio-Spin P-30 Gel Columns (Bio-Rad). The annealing was performed by boiling the oligonucleotide mix in annealing buffer (10 mM Tris-HCl (pH 8), 50 mM NaCl, 10 mM magnesium chloride) by heating for 3 min at 95 °C and slowly cooling down the mix to room temperature overnight. The sequence of the oligonucleotides used for DNA substrates preparation is listed in **Supplementary Table 2**.

### Electrophoretic mobility shift assays

DNA binding reactions (15 µl volume) were carried out in 25 mM Tris-acetate pH 7.5, 3 mM EDTA, 1 mM dithiothreitol (DTT), 100 µg/ml recombinant albumin (New England Biolabs), and 1 nM DNA substrate (in molecules). Proteins were added and reactions were incubated for 15 min on ice. Loading dye (5 µl; 50% glycerol [w/vol] and bromophenol blue) was added to the reactions and the products were separated on 6% polyacrylamide gels (ratio acrylamide:bisacrylamide 19:1, Bio-Rad) in TAE (40 mM Tris, 20 mM acetic acid and 1 mM EDTA) buffer at 4 °C. The gels were dried on 17 CHR paper (Whatman), exposed to a storage phosphor screen (GE Healthcare) and scanned by a Typhoon Phosphor Imager (FLA9500, GE Healthcare). Signals were quantified using ImageJ and plotted with GraphPad Prism 10.

## Data availability

Reagents and mouse strains are available upon request. SSDS and S1-seq data are available at Gene Expression Omnibus with the accession number xxx.

## Competing Interest Statement

The authors declare that they have no competing interests to report.

## Acknowledgments

We thank Scott Keeney for sharing *Spo11^-/-^*mice and for helpful discussions. We thank Rutgers Research Pathology Services for help with tissue processing and PAS staining. The forward genetic screen that yielded the *Mcm8^m^* allele was conducted while D.J. was a postdoctoral fellow with Scott Keeney (Memorial Sloan Kettering) and was supported by the Howard Hughes Medical Institute (to S.K) and the Human Frontier Science Program (to D.J). This work was supported by National Institutes of Health (NIH) grant R35 GM147130 (to D.J.), the National Institute of Diabetes and Digestive and Kidney Diseases intramural research program (to F.P.), the Swiss National Science Foundation (SNSF) (Grants 310030_207588 and 320030-236167) and the European Research Council (ERC) (Grant 101018257) (to P.C.). M.R.D.S. is supported by a grant from Swiss Cancer Research (KFS-6136-08-2024). Additionally, M.L. is a member of the Scott Keeney Lab (Memorial Sloan Kettering) where work is supported by the NIH grant R35 GM118092 and the Howard Hughes Medical Institute. Memorial Sloan Kettering core facilities are supported by National Cancer Institute Cancer Center support grant P30 CA08748. The contributions of the NIH author(s) are considered Works of the United States Government. The findings and conclusions presented in this paper are those of the author(s) and do not necessarily reflect the views of the NIH or the U.S. Department of Health and Human Services.

## Author Contributions

L.K.S., K.T. and M.I. performed mouse phenotyping. S.L. managed mouse husbandry. A.A. prepared the recombinant proteins and M.R.D.S. conducted *in vitro* assays. A.M.M. and N.S. performed SSDS experiments. M.L. conducted S1-sequencing. F.P. supervised A.M.M. and N.S., and P.C. supervised A.A. and M.R.D.S. D.J. and L.K.S. designed the study with contributions from all authors. D.J. and L.K.S. wrote the manuscript with contributions from all authors.

---

**Supplementary Figure 1.**
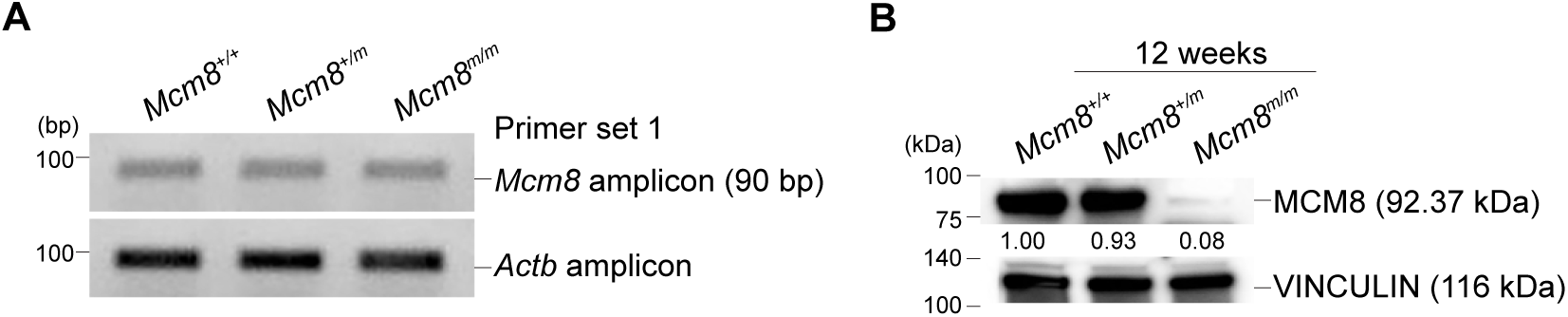
MCM8 expression analysis. **(A)** Gel electrophoresis of *Mcm8* and *Actb* RT-PCR products from adult testes (primer set 1 in Figure 2A; quantified in Figure 2C). (**B**) Immunoblot of MCM8 and VINCULIN (loading control) in adult whole-testis extracts. Relative MCM8 signal intensities normalized to VINCULIN are noted.

**Supplementary Figure 2.**
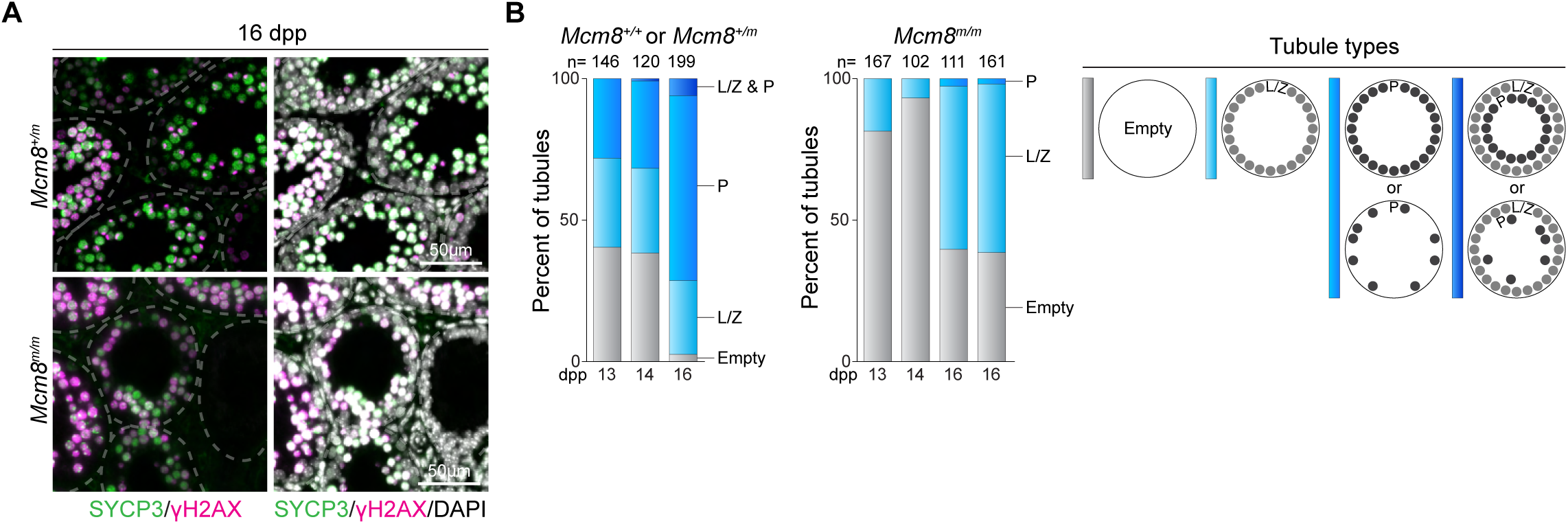
Analysis of juvenile meiosis. **(A)** Representative images of juvenile (16 dpp) seminiferous tubules immunostained for SYCP3 and γH2AX. (**B**) Bar plots show distribution of seminiferous tubule types in individual mice of the indicated ages. Tubule types (diagrammed on the right) were categorized based on presence of leptotene or zygotene (L/Z) and pachytene (P) stages, as judged by SYCP3- and γH2AX-staining patterns.

**Supplementary Figure 3.**
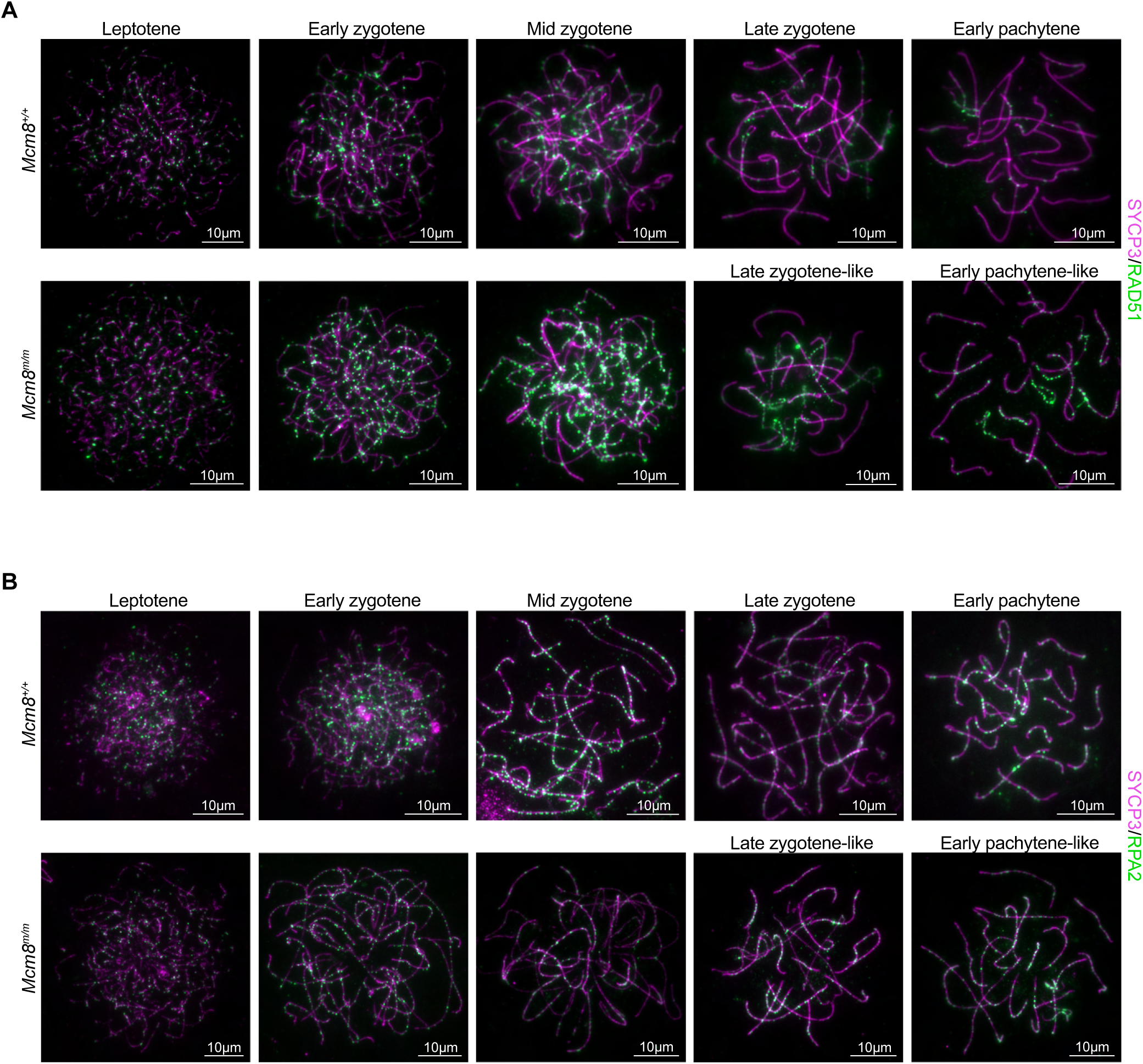
Altered recombination foci in *Mcm8* mutants. **(A)** Chromosome spreads depicting time course of RAD51 staining during meiotic prophase. (**B**) Chromosome spreads depicting time course of RPA2 staining across meiotic prophase.

**Supplementary Figure 4.**
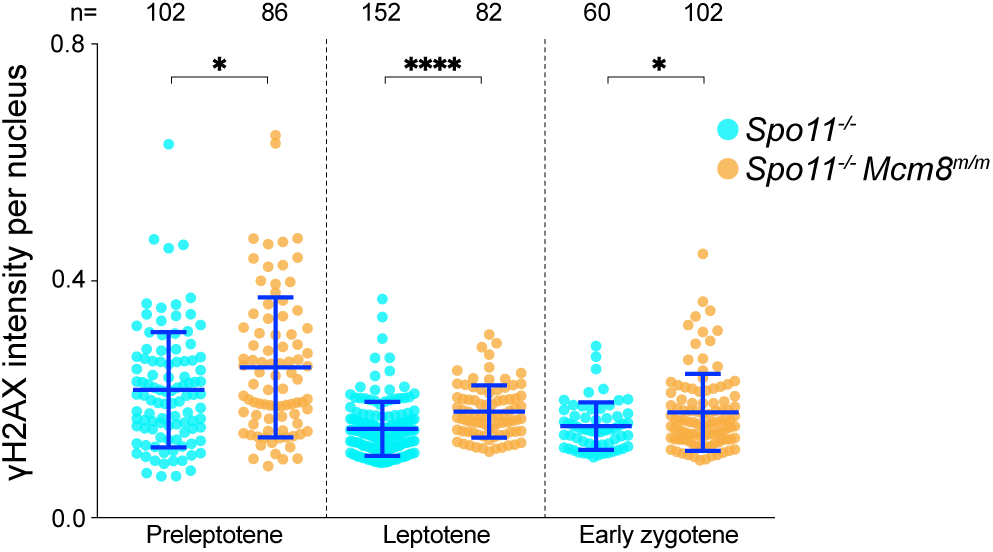
γH2AX time course in *Spo11* mutants. γH2AX intensity measurements per cell, normalized to the mean intensity of wild-type leptotene stage cells. Data from Figure 6 **(D)** was replotted with an expanded y-axis to better visualize differences between *Mcm8^+/+^* and *Mcm8^m/m^*mice.

**Supplementary Figure 5.**
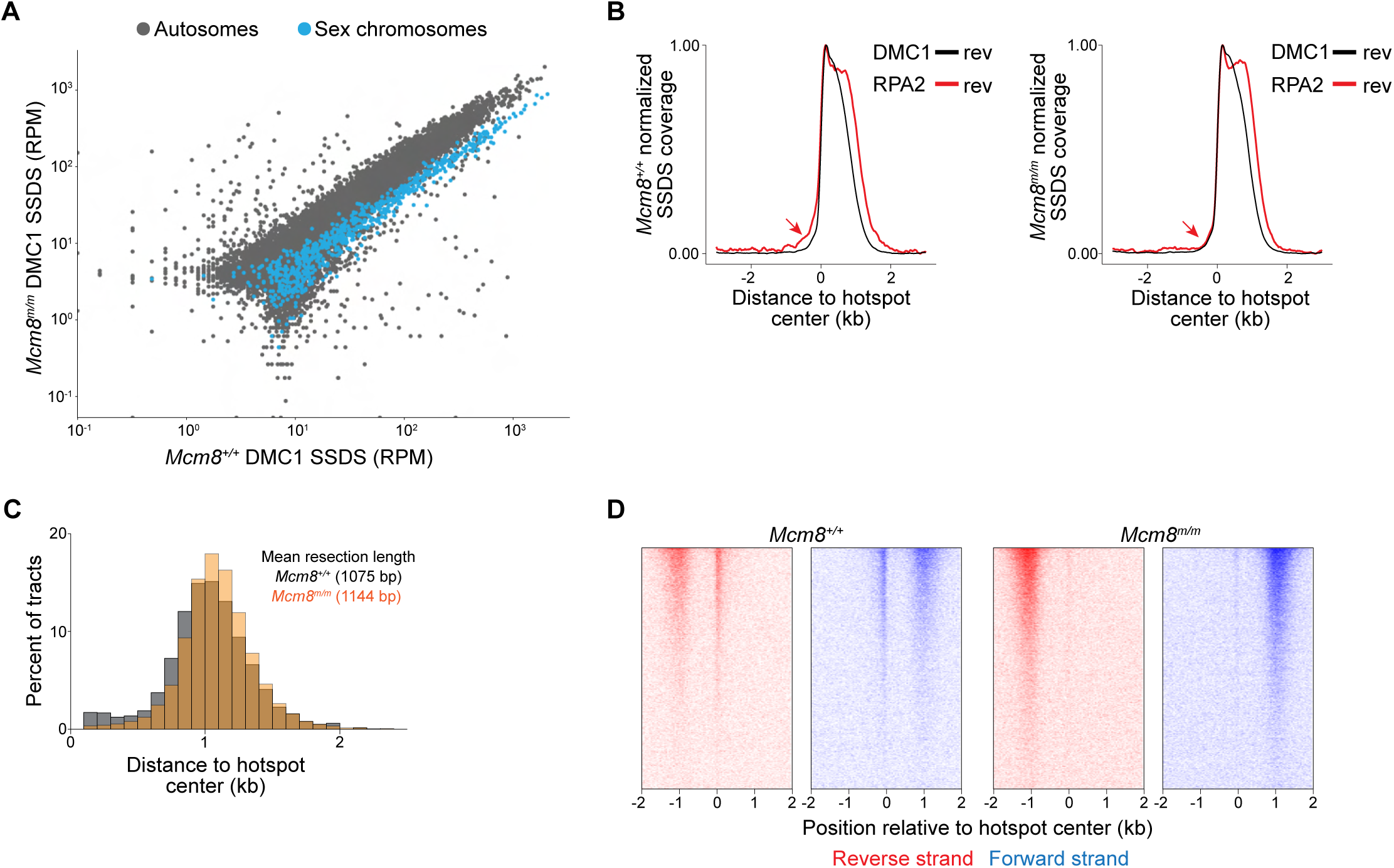
Analysis of resection in *Mcm8* mutants. (**A**) Correlation between DMC1 SSDS read counts (reads per million; RPM) at B6 hotspots present in adult *Mcm8^m/m^* and wild type-littermates. (**B**) Metaplots of DMC1 and RPA2 SSDS averages around B6 wild-type hotspots. Reads originating only from the reverse (rev) strands are shown for clarity. Red arrow in wild type points to RPA-SSDS reads that have the wrong polarity to be resection tracts. (**C**) Distribution of resection tract lengths relative to B6 wild-type hotspots. Fractions of total reads were calculated every 100 bp and plotted. Mean resection lengths are indicated. (**D**) Heatmaps of strand-specific S1-seq reads counts around B6 wild-type hotspots in adult mice. Each row is a single hotspot, ranked from the strongest at the top.

**Supplementary Figure 6.**
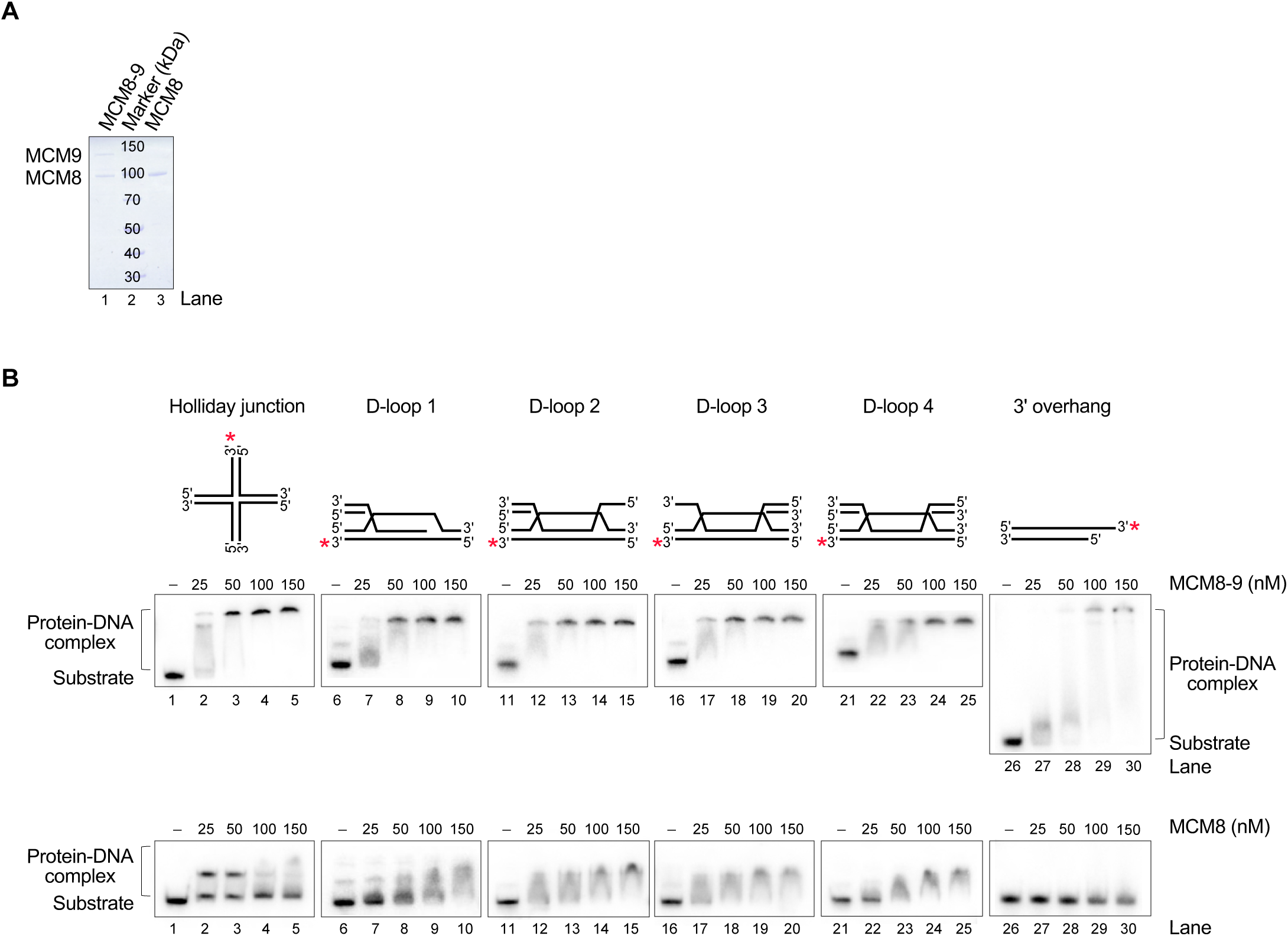
MCM8 binding to oligonucleotide-based DNA structures. (**A**) Recombinant MCM8-9 complex and MCM8 alone used for the biochemical experiments. The polyacrylamide gel was stained with Coomassie Brilliant Blue. (**B**) Representative gels for binding assays (quantified in Figure 8). Schematics of substrates used are shown on the top and red asterisks represent the positions of the radioactive labels.

**Supplementary Table 1.** Antibodies.

| **Primary Antibody** | **Company** | **Reference** | **Host Species** | **Application** | **Concentration** |
| --- | --- | --- | --- | --- | --- |
| DMC1 | ProteinTech | 13714-1-AP | Rabbit | Spread | 1:100 |
| DMC1 | Gift from Florencia Pratto |  | Rabbit | Spread; SSDS | 1:50; 2 µg |
| γH2AX | Novus Biological | NB100-384 | Rabbit | Spread; Histology | 1:1000; 1:250 |
| MSH5 | Gift from Florencia Pratto |  | Rabbit | Spread | 1:50 |
| MLH1 | Cell Signaling | 3515S | Mouse | Spread | 1:25 |
| MCM8 | ProteinTech | 16451-1-AP | Rabbit | Section; WB | 1:100; 1:2500 |
| RAD51 | Millipore-Sigma | pc130 | Rabbit | Spread | 1:100 |
| RPA2 | Abcam | ab10359 | Mouse | SSDS | 2 µg |
| RPA2 | Cell Signaling | 2208S | Rat | Spread | 1:75 |
| STRA8 | Abcam | ab49602 | Rabbit | Histology | 1:1000 |
| SYCP1 | Abcam | ab15090 | Rabbit | Spread | 1:100 |
| SYCP3 | Novus Biological | NB300-232 | Rabbit | Spread | 1:200 |
| SYCP3 | Abcam | ab15093 | Rabbit | Spread | 1:200 |
| SYCP3 | Santa Cruz | sc-74569 | Mouse | Spread; Histology | 1:200; 1:250 |
| VINCULIN | Santa Cruz | sc-73614 | Mouse | WB | 1:2500 |
| **Secondary Antibody** | **Company** | **Reference** | **Host Species** | **Application** | **Concentration** |
| Mouse 488 | Invitrogen | A21202 | Donkey | Spread; Histology | 1:250; 1:500 |
| Mouse 594 | Invitrogen | A11032 | Goat | Spread; Histology | 1:250; 1:500 |
| Mouse Digital | Kindle Biosciences | R1005 |  | WB | 1:2000 |
| Rabbit 488 | Invitrogen | A11034 | Goat | Spread; Histology | 1:250; 1:500 |
| Rabbit 594 | Invitrogen | A21207 | Donkey | Spread; Histology | 1:250; 1:500 |
| Rabbit Digital | Kindle Biosciences | R1006 |  | WB | 1:1000 |
| Rat 488 | Invitrogen | A11006 | Goat | Spread | 1:250 |

**Supplementary Table 2.**
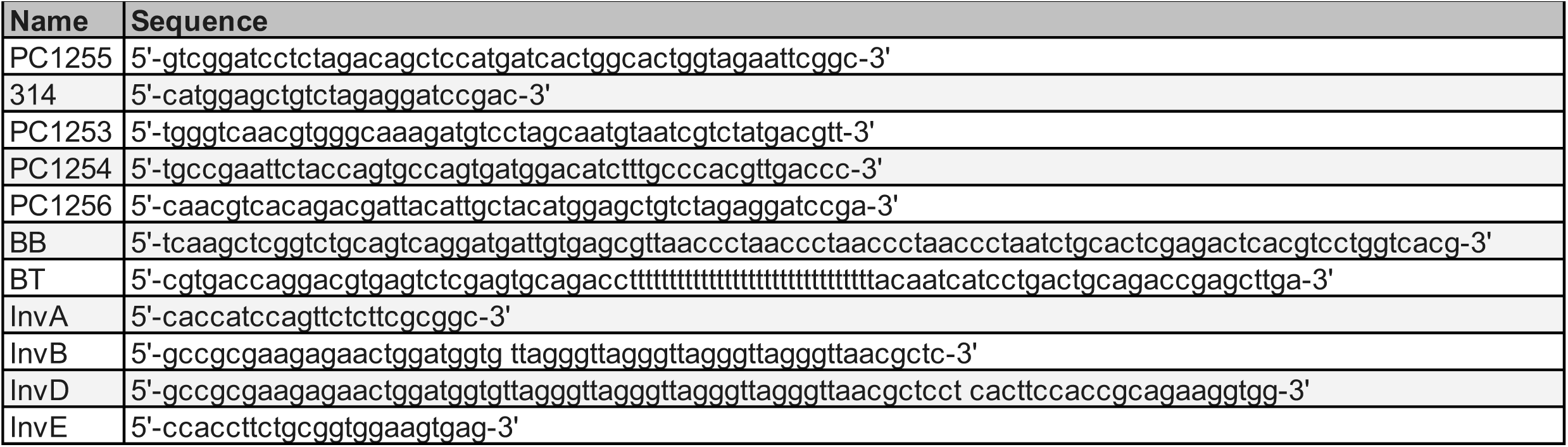
Oligonucleotides used for preparation of DNA substrates.

## Notes

### Competing Interest Statement

The authors have declared no competing interest.

### Summary of Updates

This version of the manuscript has been revised to update the acknowledgments section. Remainder of the manuscript is unchanged.

